# Cyclin O controls entry into the cell-cycle variant required for multiciliated cell differentiation

**DOI:** 10.1101/2024.05.22.595363

**Authors:** Michella Khoury Damaa, Jacques Serizay, Rémi Balagué, Amélie-Rose Boudjema, Marion Faucourt, Nathalie Delgehyr, Kim Jee Goh, Hao Lu, Ee Kim Tan, Cameron T. James, Catherine Faucon, Rana Mitri, Diana Carolin Bracht, Colin D. Bingle, Norris Ray Dunn, Sebastian J. Arnold, Laure-Emmanuelle Zaragosi, Pascal Barbry, Romain Koszul, Heymut Omran, Gabriel Gil-Gόmez, Estelle Escudier, Marie Legendre, Sudipto Roy, Nathalie Spassky, Alice Meunier

## Abstract

Multiciliated cells (MCC) ensure proper fluid circulation in various organs in metazoans. Their differentiation is marked by the massive ampliication of cilia-nucleating centrioles and is known to be controlled by various cell cycle components. Tn a companion study, we show that the differentiation of MCC is driven by a genuine cell-cycle variant characterized by sequential and wave-like expression of canonical and non-canonical cyclins such as Cyclin O (CCNO). Patients with *CCNO* mutations exhibit a subtype of Primary Ciliary Dyskinesia (PCD) designated as Reduced Generation of Multiple Motile Cilia (RGMC), yet the role of CCNO during MCC differentiation remains unclear. Here, using mice and human cellular models, single cell transcriptomics and functional studies, we show that *Cena* is activated during a strategic temporal window at the crossroads between the onset of MCC differentiation, the entry into the MCC cell cycle variant, and the activation of the centriole biogenesis program. We ind that the absence of *Cena* leads to a block of MCC progenitor differentiation at the G1/S-like transition, just before the beginning of centriole formation. This leads to a complete lack of centrioles and cilia in mouse brain and human airway MCC. Altogether, our study identifies CCNO as a core regulator of entry into the MCC cell cycle variant and shows that the coupling of centriole biogenesis to an S-like phase, maintained in MCC, is dependent on CCNO.

**One sentence summary:** Cyclin O is necessary for multiciliated cells to enter their differentiation cell cycle variant and allows the massive amplification of centrioles, which serve as basal bodies for cilia nucleation.

## Introduction

Multiciliated cells (MCC) propel physiological fluids within the lumen of several fluid-producing organs. Tn the brain ventricles, they contribute to the flow of cerebrospinal fluid; in the respiratory tract, they are necessary for mucociliary clearance; and in the female and male reproductive tracts, they contribute to the movement of eggs and sperm, respectively (Aprea et al., 2021; Spassky and Meunier, 2017; Terre et al., 2019; Yuan et al., 2019). A key event during MCC differentiation is the massive ampliication of centrioles, which then mature as basal bodies for nucleating tufts of motile cilia. Although MCC from different tissues do not share the same cell lineage, the molecular cascade that regulates centriole amplification seems well conserved and shares similarities with the centriole duplication program during the cell cycle. This process requires critical centriolar assembly proteins such as SAS6 and PLK4 (LoMastro et al., 2022; Vladar and Stearns, 2007; Zhao et al., 2013) and master cell cycle regulators such as CDK1, CDK2, PLK1 and APC/C that control centriole number and maturation (Al Jord et al., 2017; Kim et al., 2022; Vladar et al., 2018). Tn a companion study, we further show that these regulators constitute the tip of the iceberg and that, in fact, an entire cell cycle variant, re-expressing the majority of cell cycle players, progressing through S-, G2- and M-like phases and tailored with wave-like expression of canonical and non-canonical cyclins, is at work during MCC differentiation. This cell cycle variant also seems conserved in various mouse tissues and in human airways (Serizay et al., 2024).

The contribution of Cyclin O (CCNO) -the first non-canonical cyclin expressed during this cell cycle variant- to MCC differentiation was first discovered in human patients with mutations in the *CCNO* gene. These patients suffer from a distinctive form of Primary Ciliary Dyskinesia (PCD) referred to as Reduced Generation of Multiple Motile cilia (RGMC) because of cilia rarity in the airway epithelium (Amirav et al., 2016; Casey et al., 2015; Guo et al., 2020; Henriques et al., 2021; Ma et al., 2021; Wallmeier et al., 2014). Tn parallel, an in-depth study of airway MCC from *Cena* mutant mice showed a disorganized ampliication of centrioles, ultimately resulting in incomplete multiciliation (Funk et al., 2015; *Cena^RA^*). Similarly, in the multiciliated epidermis of Xenopus larvae -a surrogate model of mammalian MCC-bearing epithelia-morpholino-mediated inhibition of CCNO function results in a reduced number of centrioles and cilia (Amirav et al., 2016; Wallmeier et al., 2014). Although not comprehensively studied at the cellular level, the phenotype appears similar in the oviducts of full *Cena* knockout mice, with a decrease in the number of cilia per cell (N(nez-Olle et al., 2017), and in the efferent ducts of the testes, where a loss of multiciliation is observed (Terre et al., 2019). Tn mouse brain, the cellular phenotype appears more severe but is not quantified (N(nez-Olle et al., 2017).

The clinical phenotype of RGMC patients with *CCNO* mutations unequivocally deines CCNO as a determinant of MCC function. However, the cellular phenotype of CCNO depletion varies from a delayed and defective centriole amplification leading to partial multiciliation, to a total absence of cilia, depending on how CCNO is depleted, the cellular model and the species examined (Funk et al., 2015; N(nez-Olle et al., 2017; Terre et al., 2019; Wallmeier et al., 2014). Moreover, the mechanism of action of CCNO in the MCC differentiation program remains unknown. Here, using two mouse models for *Cena* loss-of-function, as well as airway MCC derived from human embryonic cells (hESCs) and nasal biopsies from PCD patients with *CCNO* mutations, we show that (i) a subpopulation of airway MCC from *Cena* mutant mice can amplify centrioles, albeit abnormally, while human airway MCC cannot, that (ii) mouse brain MCC display a phenotype comparable to that of human airway MCC, where centrioles are not produced in the absence of CCNO and that (iii) CCNO controls centriole amplification by regulating entry into the MCC differentiation cell-cycle variant.

## Results

### Mutated *Ccno* progenitors fail to develop into MCC in the mouse brain compared to the mouse trachea

Discrepancies between cellular models in the literature prompted us to test whether a differential phenotype could exist between brain and airway MCC from mice without CCNO. Human and mouse CCNO proteins share similarities with other canonical cyclins such as CCNA, CCNB and CCNE (**Fig. S1A**). The *Cena* genetic locus is overall well conserved across vertebrates, exhibiting collinearity with *Meidas* immediately downstream of *Cena,* although we note that *Cena* and *Meidas* collinearity is absent in Zebrafish (Defosset et al., 2021, **Fig. S1A**). We first leveraged the *Cena^RA^* mouse model, where 2 out of 3 exons of *Cena* are lacking (**Fig. S1B,** Funk et al., 2015), and performed whole tissue immunostainings of glutamylated tubulin (GT335) to detect multiple cilia of MCC, and the primary cilium of other cell types including MCC progenitors, from samples of both lateral ventricular walls of the brain and trachea. To circumvent individual variability, we compared brain and airway epithelia dissected from the same individuals at postnatal day 35 (P35), when MCC differentiation is complete in both tissues. As previously published (Funk et al., 2015), we observed MCC in *Cena^RA^* tracheal tissues, albeit more scarcely distributed than in the controls. However, in the same individuals, MCC are very rarely observed in brain tissues, while they cover the whole epithelium in the brains of control animals (**Fig. S1C**).

To test whether the truncated CCNO protein that could potentially be translated in the *Cena^RA^* model could be responsible for the partial multiciliation observed in tracheal tissues, we then leveraged mutants with a complete deletion of the *Cena* gene locus (*Cena^KO^*, **Fig. S1B**, N(nez-Olle et al., 2017). Here again, almost no MCC were observed in the brain, while in the trachea of the same individuals, MCC can still be formed, albeit with approximately 70% reduction compared to controls (**Fig. 1A and B**). Tn the mutants’ brain, the few MCC produced have an abnormal morphology, with very few multicilia. Tn the trachea though, half of them present a seemingly normal morphology while the other half present more scarcely distributed multicilia, as previously described in mouse *Cena^RA^*respiratory cells (**Fig. 1C and D;** Funk et al., 2015).

**Figure 1.**
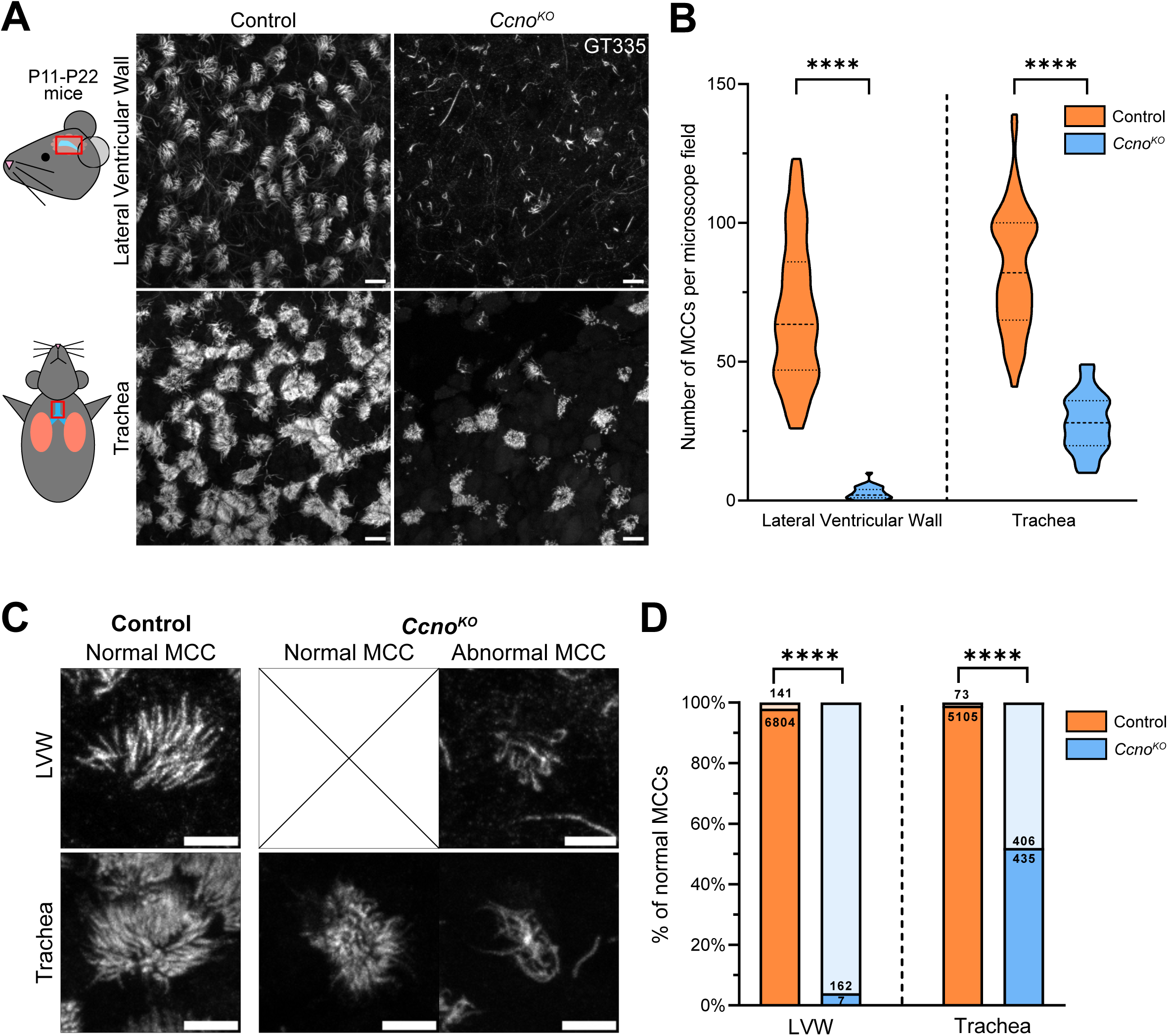
Mutated *Ccno* progenitors fail to develop into MCC in the mouse brain compared to the mouse trachea. **(A)** Tmmunostaining of control and *Cena^KO^* mice lateral ventricular wall (LVW) of the brain and trachea dissected from the same animal at P22, with polyglutamylation marker GT335 for cilia. Scale bar 10µm. **(B)** Quantification of MCC per microscope field in both LVW and trachea of P11 to P22 mice. Tmages from 12 control mice and 6 *Cena^KO^* mice were quantified, with 3 to 6 images per immunostained tissue, Control LVW: 100 values, *Cena^KO^* LVW: 60 values, Control trachea: 59 values, *Cena^KO^* trachea: 30 values. P-values derived from two-tailed Mann-Whitney U-test, ****p-value<0.0001. **(C)** The few MCC found in the brain are almost all abnormal in the *Cena^KO^*, with very few and disorganized cilia per cell, as shown by upper panels. Tn the *Cena^KO^* trachea, half of the formed MCC have a normal morphology; the rest is abnormal as indicated by the bottom panels. Scale bar 5µm. **(D)** Quantification of normal MCC in both LVW and trachea of P11 to P22 mice. P-values derived from two-sided Chi-square test (two-proportion z-test), ****p-value<0.0001.

Altogether, this shows that in both *Cena* mutants, a differential phenotype exists between brain and tracheal MCC which suggests that in mouse, CCNO is essential for MCC formation in the brain while its absence can be partially compensated for in the tracheal tissues. Due to the wide range of severe clinical manifestations of patients with *CCNO* mutations (Amirav et al., 2016), we decided to further assess the function of CCNO in MCC differentiation, using mouse brain MCCs as a model system.

### CCNO stands at the crossroads between MCC differentiation, MCC cell cycle variant and centriole amplification pathways

We sought to identify the molecular and cellular processes of MCC differentiation that are controlled by CCNO. We first performed immunostaining for CCNO protein during in vitro differentiation of MCC brain progenitors. We found that CCNO protein is absent before the onset of centriole ampliication, when progenitors display a single centrosome (labelled by FOP). CCNO is then detected as cells enter the amplification “A’-phase, when procentrioles amplify, and during the growth “G’-phase, when procentrioles mature, with enrichment in the nuclear compartment. Eventually, CCNO is sharply reduced in both cell compartments during the disengagement “D’-phase, when centrioles begin to disengage from their growing platforms, the deuterosomes, and until multiciliation (**Fig. 2A and B, Fig. S2,** Al Jord et al., 2017). CCNO seems to be mostly nuclear located as shown by intensity ratio of nuclear/cytoplasmic signal in A and G-phases (**Fig. 2B**).

**Figure 2.**
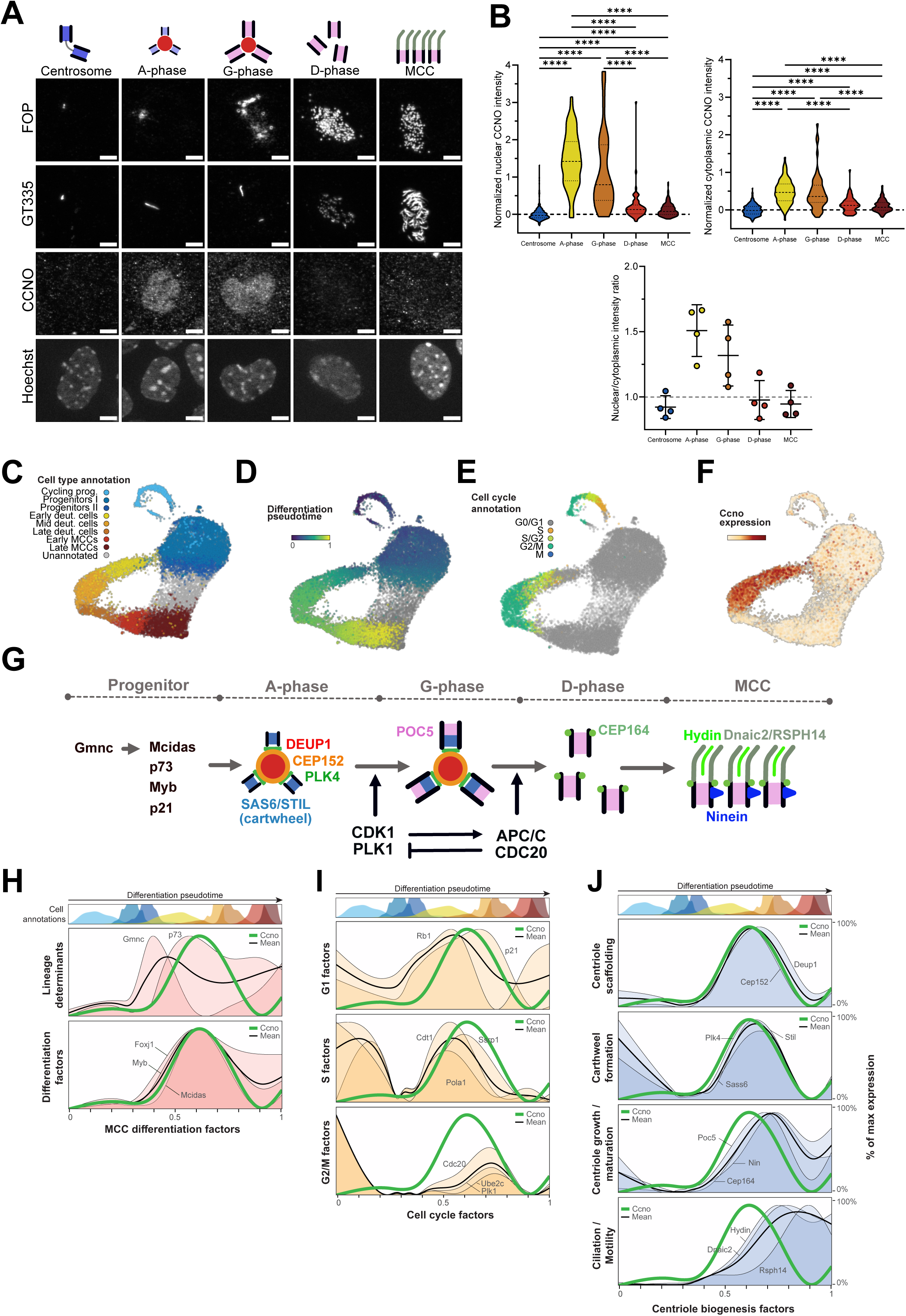
Cyclin O is expressed at the onset of MCC differentiation, MCC ell cycle variant and MCC centriole amplification. **(A)** Tmmunostaining of ependymal cells in culture at Day-Tn-Vitro 5 (DTV5) for FOP (centrioles), GT335 (cilia) and CCNO. CCNO staining is shown for the main stages of centriolar multiplication, from the progenitor 2-centrioles state to the fully mature multiciliated cell (MCC). Centrioles are represented by rectangles with black borders, deuterosomes by red circles and cilia by grey lines. Scale bar 5µm. **(B)** Nuclear and cytoplasmic intensity quantification of CCNO staining during the different stages shown in (A), normalized to the mean of centrosome stage intensity. Bottom graph shows mean nuclear/cytoplasmic intensity ratio per quantified coverslip for each stage. n=3. Nuclear intensity: Centrosome: 470 cells, A-phase: 92 cells, G-phase: 59 cells, D-phase: 66 cells, MCC: 188 cells. Cytoplasmic intensity: Centrosome: 269 cells, A-phase: 92 cells, G-phase: 58 cells, D-phase: 63 cells, MCC: 175 cells. P-values are derived from Kruskal-Wallis test + Dunn’s multiple comparisons, ****p-value<0.0001. **(C)** Single-cell RNA-seq data of cultured ependymal cells at DTV2 and their cluster annotation in UMAP projection, as described in Serizay et al., 2024. **(D)** UMAP projection of cells coloured by their pseudotime (Serizay et al., 2024), from 0 (cycling progenitors) to 1 (late MCC) with intermediate values attributed to progenitors and differentiating cells. **(E)** UMAP projection of cells coloured by their putative cell cycle phase annotations inferred from neural stem cell reference (O’Connor et al., 2021). **(F)** UMAP projection of cells coloured by their *Cena* level of expression. Expression spans early and mid deuterosomal cells clusters described in (C). **(G)** Schematic representation of the main differentiation actors and centriolar assembly proteins involved in MCC differentiation steps. **(H)** Expression of several MCC differentiation factors along differentiation pseudotime and their mean expression (black line), compared to *Cena* expression (green thick line). **(I)** Expression of cell cycle factors along differentiation pseudotime and their mean expression (black line), compared to *Cena* expression (green thick line). **(J)** Expression of centriole biogenesis factors along differentiation pseudotime and their mean expression (black line), compared to *Cena* expression (green thick line).

To further reine the context in which *Cena* is expressed, we leveraged the MCC differentiation lineage pseudotime inferred from the single cell transcriptomics dataset described in Serizay et al., 2024, comprising (i) cycling and quiescent brain MCC progenitors, (ii) differentiating MCC progenitors (identified as early, mid and late deuterosomal cells) and (iii) terminally differentiated MCC (**Fig. 2C-E**). Tn this continuous transcriptional landscape and consistent with immunostaining data, *Cena* is not expressed in MCC progenitors, is activated in the early and mid deuterosomal cell clusters and is then silenced along the rest of the differentiation process (**Fig. 2C-F**). Cell cycle phase annotations from a neural stem cell single-cell RNA-seq reference (O’Connor et al., 2021) detects *Cena* expressing cells as progressing from G0/G1 to S/G2 and G2/M-like phases of the cell cycle (**Fig. 2E**). Finally, we show in our associated study (Serizay et al., 2024) that *Cena* expression is temporally correlated with the expression of centriole biogenesis core components.

We further analysed the temporality of *Cena* expression compared to core regulators of (i) MCC fate determination, (ii) MCC cell cycle variant and (iii) centriole biogenesis. First, the temporality of expression of MCC fate determinants is consistent with functional studies (Lewis and Stracker, 2020; Lyu et al., 2024) where the lineage determinant *Gmne* is expressed first, specifically in the progenitor and early deuterosomal clusters, followed by *Myb*, involved in multilineage airway epithelial cell differentiation (Pan et al., 2014) expressed from late progenitor to mid-deuterosomal clusters. *Fax}1* is also upregulated in the progenitor clusters but remains expressed during the whole differentiation process, consistent with its role in late events during MCC differentiation. Finally, transcription factor *p73* expression occurs early on in the early deuterosomal cluster, whereas *Meidas* is expressed slightly later, in early and mid-deuterosomal clusters (**Fig. 2G-H, Fig. S3A**). Tn this transcriptional landscape and consistent with identifying *Cena* as a Myb-regulated gene (Pan et al., 2014), *Cena* activation and duration of expression is comparable to *Myb* from late progenitor to mid-deuterosomal clusters. Unexpectedly though, *Cena* activation seems to occur slightly before *Meidas.* (**Fig. 2G-H, Fig. S3A**). Within the MCC cell cycle variant, and consistent with cell cycle phase annotations, the onset of *Cena* expression is consecutive to the activation of typical G1 factors (*Cdkn1A* (*p21*) and *Rb1*), is concomitant with S factors (*Cdt1*, FACT complex subunit *Ssrp1* and the catalytic subunit of DNA polymerase alpha (*Pala1*)), and precedes and overlaps the activation of factors involved in mitotic progression (*Plk1*, *Ube2e* subunit of APC/C, APC/C cofactor *Cde20*). These cell cycle core regulators are silenced before or, at last, concomitantly with *Cena* (**Fig. 2I**).

Finally, concerning the centriole biogenesis molecular subprocesses, *Cena* activation is concomitant with the centriole scaffolding genes *Deup1* and *Cep152,* and slightly precedes the expression of cartwheel components *Plk4*, *Stil* and *Sass6.* The duration of expression of all these early genes is comparable with *Cena*. The onset of expression of genes involved in centriole growth and maturation (*Pae5*, *Cep164* and *Ninein*) occurs later, and continues after *Cena* silencing. Finally, the motile cilia genes *Dnai2*, *Hydin and Rsph14* are expressed later, only partially overlapping with decreasing expression of *Cena* (**Fig. 2J, Fig. S3B**).

Together, these observations highlight the strategic temporal window during which *Cena* is activated, at the crossroads between the onset of MCC differentiation, the entry into the MCC cell cycle variant and the activation of the centriole biogenesis program.

### CCNO is required for the early progression of MCC differentiation

We sought to decipher at which step of MCC differentiation CCNO is required. We performed single-cell RNA-seq on differentiating ependymal progenitor cells harvested *in-vitra* from three different *Cena^KO^* individuals at DTV2 (**see methods**) and compared the cell composition with that observed in scRNAseq from wildtype (WT) individuals (**Fig. S4A**). Tn *Cena^KO^* mutants, cells at the mid- and late deuterosome stages are no longer observed, indicating that the MCC cell differentiation process is aborted early on. Consistent with this inding, a drastic reduction of terminally differentiated MCC is observed, recapitulating immunostaining observations. By contrast and as expected, proliferating progenitors, which do not express *Cena*, are unaffected in *Cena^KO^*samples (**Fig. S4A**).

We inferred a shared differentiation lineage from co-embedded WT and *Cena^KO^* cells. We used it to precisely identify the lineage timepoint at which *Cena^KO^* deuterosomal cells stop differentiating (**Fig. S4B**, **S4C, methods**), corresponding to a transcriptional state reached by deuterosomal cells in both WT and *Cena^KO^* conditions. We re-annotated deuterosomal cells prior to this lineage timepoint as **primordial deuterosomal cells** (**Fig. 3A**). Tn *Cena^KO^*primordial deuterosomal cells, the master regulator of MCC differentiation *Gmne* is not differentially expressed, confirming that *Cena^KO^* cells successfully engage into the MCC fate (**Fig. 3B**). Consistently, *Faxj1, Myb and p73* are also not differentially expressed in *Cena^KO^* primordial deuterosomal cell*s* (**Fig. 3B, Fig. S5A**), and FOXJ1+ cells are present in comparable proportions in *in vitra* culture of WT or *Cena^KO^* differentiating radial glial mouse progenitors (**Fig. 3C**). Tn contrast, *Meidas* expression is halved in *Cena^KO^* primordial deuterosomal cells compared to their WT counterparts (**Fig. 3B**), contrary to what has been previously documented in mTECs by bulk RT-qPCR (Funk et al., 2015). We transfected *Cena^KO^* cells with *Meidas* or *Meidas + Cena* and could only restore multiciliogenesis when both factors were re-expressed, indicating that the MCC differentiation arrest in *Cena^KO^* is not caused by the decreased expression of *Meidas* (**Fig. S5B**). To further characterize the transcriptional landscape preceding the differentiation arrest, we performed genome-wide differential expression analysis between WT and *Cena^KO^* primordial deuterosomal cells (**Fig. 3D**). We identified dozens of differentially expressed genes such as *Cde20, Tap2a, Aurka, Wee1 and Plk4* genes that are down-regulated in *Cena^KO^*cells, or *Cfap53*, *Pifa* and *Spag1* genes that are up-regulated in *Cena^KO^* cells. Using genome-wide gene set enrichment analysis (GSEA), we found that sets of genes involved in cell cycle pathways, including cell cycle regulation and centrosome regulation, are overall down-regulated in *Cena^KO^*primordial deuterosomal cells. Tn contrast, sets of genes involved in cilia biogenesis and motility are up-regulated in *Cena^KO^* primordial deuterosomal cells (**Fig. 3E**). These results reveal that in absence of *Cena,* MCC progenitors can enter the MCC differentiation lineage and even trigger the expression of cilia biogenesis factors. However, they arrest early on in the differentiation process, concomitantly with a decreased expression of cell cycle and centriole biogenesis factors.

**Figure 3.**
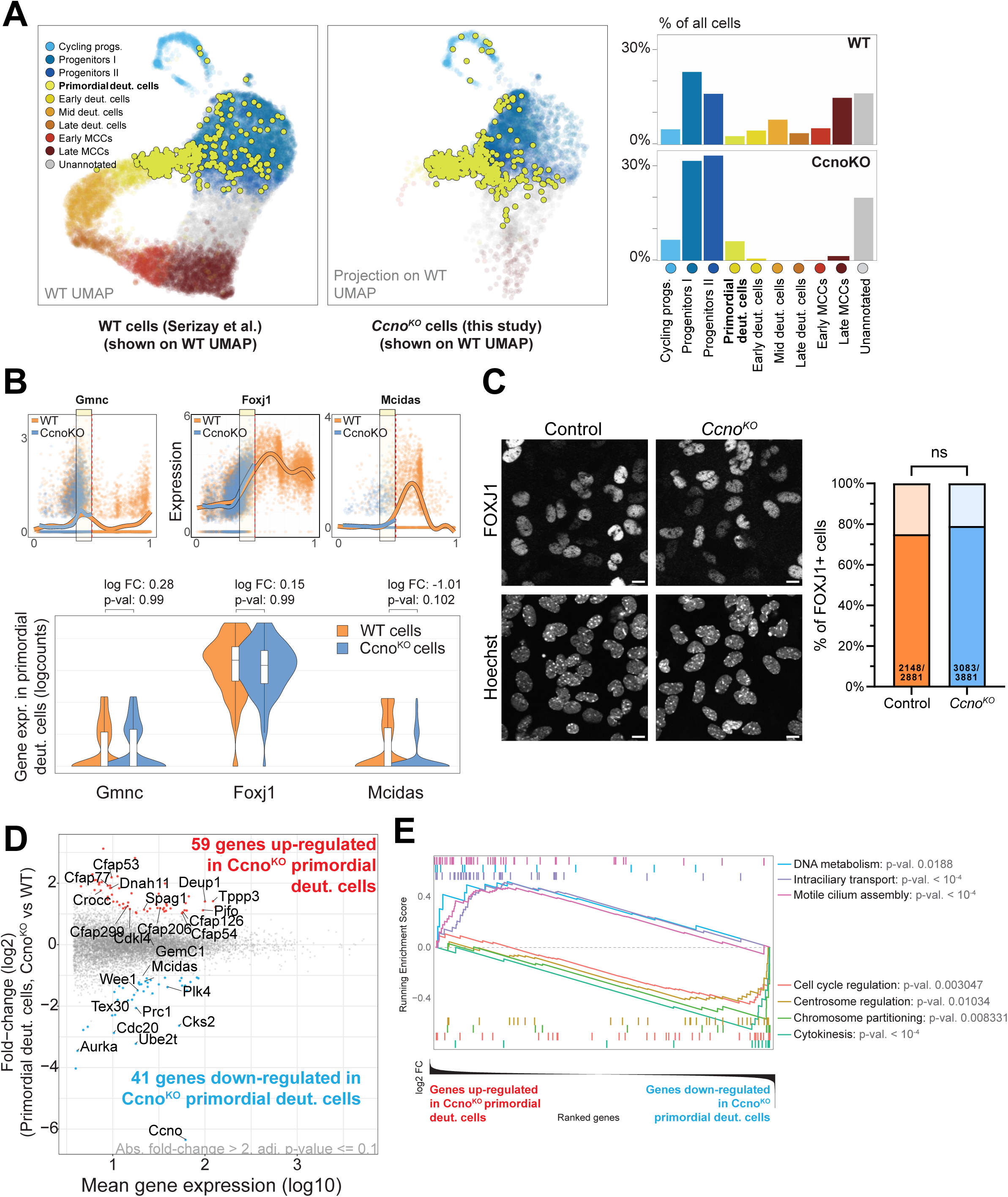
CCNO is required for the early progression of MCC differentiation. **(A)** scRNAseq data of *Cena^KO^* cells compared to the previously shown data set of WT cells (Fig. 2C). *Cena^KO^*cells are projected onto the WT UMAP embedding and annotations are transferred from WT to *Cena^KO^* cells. Note that very few cells in the deuterosomal clusters are present in the *Cena^KO^*. Primordial deuterosomal cells cluster, present in WT and *Cena^KO^* samples, are shown in yellow. **(B)** MCC differentiation factors *Gmne, Fax}1* and *Meidas* are expressed in primordial deuterosomal cells from WT and *Cena^KO^* samples. *Gmne* and *Fax}1* expression in primordial deuterosomal cells is similar in WT and *Cena^KO^*, and *Meidas* expression is slightly downregulated in the *Cena^KO^* primordial cells. **C)** Left panel: Tmmunostaining of FOXJ1 in in-vitro ependymal cells cultures. Scale bar 10µm. Right panel: Quantification of the proportion of control vs *Cena^KO^* FOXJ1 positive cells in cultures from DTV2 to DTV23, showing that FOXJ1 expression is not affected in *Cena^KO^* cells at the protein level. P-values derived from two-sided Chi-square test (two-proportion z-test), ns: not signiicant. **(D)** Genome-wide differential expression analysis between WT and *Cena^KO^* primordial deuterosomal cells cluster, showing 59 genes upregulated in the *Cena^KO^* and 41 downregulated genes. **(E)** Gene set enrichment analysis (GSEA) of genes ranked by their expression change in *Cena^KO^* versus WT primordial deuterosomal cells, showing upregulation of genes involved in cilia motility in *Cena^KO^* primordial cells and a downregulation of genes involved in cell cycle and centrosome regulation.

### Absence of CCNO blocks the progression through the MCC cell cycle variant

Waves of canonical and non-canonical cyclins segment the MCC cell cycle variant, described in the companion study, in successive phases. Cyclin O is the main Cyclin expressed during the first part of the MCC cell cycle variant (Serizay et al., 2024). We further analysed the *Cena^KO^* scRNAseq data and found that, whereas WT deuterosomal cells can progress into S-like and G2/M-like phases, *Cena^KO^* primordial deuterosomal cells arrest just before the transition into the S-like phase (**Fig. 4A and B**). Consistent with CCNO being phylogenetically closer to canonical cell cycle cyclins than to atypical cyclins (Quandt et al., 2020, **Fig. S1A**), this suggests that CCNO is required for the progression of progenitor cells through cell cycle-like phases of differentiation, comparable to the role of the canonical CCNE2 for the progression through the G1/S phase of the canonical cell cycle.

**Figure 4.**
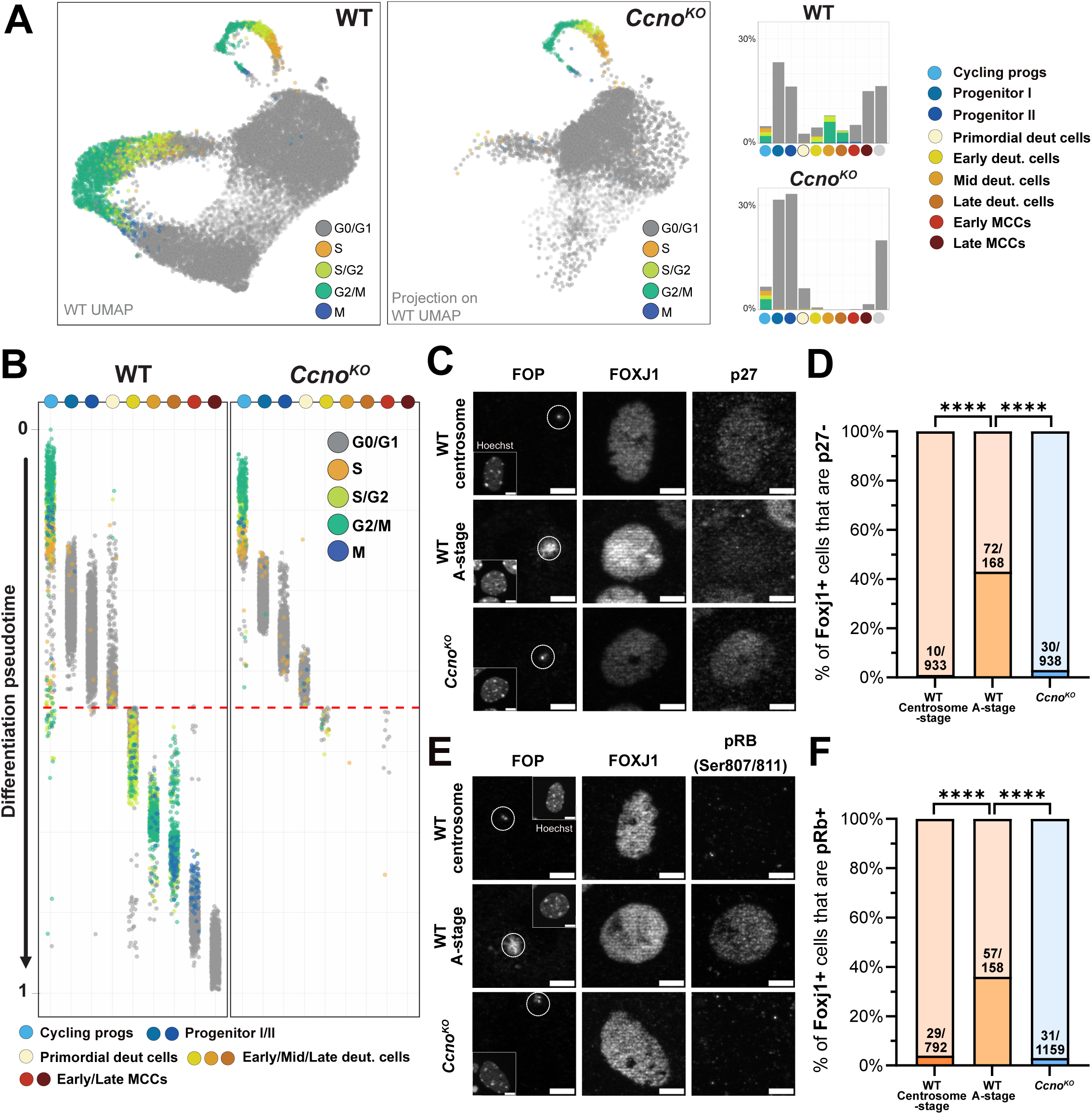
Absence of CCNO blocks the progression through the MCC cell cycle variant. **(A)** Putative cell cycle phase annotations of WT and *Cena^KO^* cells, showing a lack of S/G2/M-like deuterosomal cells in the *Cena^KO^*. **(B)** Putative cell cycle phase annotations for each cluster in WT and *Cena^KO^*samples along a shared differentiation pseudotime, showing that *Cena^KO^* are blocked at the G0/G1-like phase before entry in S-like phase. **(C)** Tmmunostaining of in-vitro WT and *Cena^KO^* MCC progenitors at DTV5 for FOP, Foxj1 and p27, typically degraded in canonical S-phase, and during MCC differentiation from A-stage. Centrioles of the centrosome are FOP+. A-stage of centriole amplification is identified by the formation of a FOP+ cloud. Cells with a FOP+ cloud are not observed in *Cena^KO^.* Scale bar 5µm. **(D)** Quantification of p27-cells in FOXJ1+ WT centrosome stage cells, FOXJ1+ WT A-stage cells and FOXJ1+ *Cena^KO^* cells. Tn FOXJ1+ cells, *Cena^KO^* cells do not degrade p27 compared to FOXJ1+ WT cells that are in the A-stage and are similar to FOXJ1+ WT cells that are in the centrosome stage, just before centriole amplification. n=3. P-values derived from two-sided Chi-square test (two-proportion z-test), ****p-value<0.0001 **(E)** Tmmunostaining of in-vitro WT and *Cena^KO^*MCC progenitors at DTV5 for FOP, FOXJ1 and pRB (Ser807/Ser811), classically phosphorylated in S-phase, and during MCC differentiation from A-stage. Centrioles of the centrosome are FOP+. A-stage of centriole amplification is identified by the formation of a FOP+ cloud. Cells with a FOP+ cloud are not observed in *Cena^KO^.* Scale bar 5µm. **(F)** Quantification of pRB+ cells in FOXJ1+ WT centrosome stage cells, FOXJ1+ WT A-stage cells and FOXJ1+ *Cena^KO^* cells. Tn FOXJ1+ cells, *Cena^KO^* cells do not phosphorylate RB compared to FOXJ1+ WT cells that are in the A-stage and are similar to FOXJ1+ WT cells that are in the centrosome stage, just before centriole amplification. n=3. P-values derived from two-sided Chi-square test (two-proportion z-test), ****p-value<0.0001.

To validate the role of *Cena* for progression through successive phases of the MCC cell cycle variant, we performed immunostainings on p27^Kip1^, whose degradation is a hallmark of cycling cells entering S phase (Coats et al., 1996) and of differentiating MCC progressing through the A-, G- or D-stages of centriole ampliication (Al Jord et al., 2017). We quantified p27 negativity in FOXJ1+ WT or *Cena^KO^* cells. Tn contrast with the increasing number of FOXJ1+/p27-cells in WT progressing into the MCC differentiation early A-stage (“= 40%), we observed very few FOXJ1+/p27-cells in FOXJ1+ *Cena^KO^* cells (“= 4%), more close to WT FOXJ1+ cells before centriole ampliication, at the “centrosome’ stage (“= 1%, **Fig. 4C and D**). We also stained WT and *Cena^KO^* cultures for retinoblastoma protein phosphorylation (pRB pSer807/811), another hallmark of cycling cells entering S-phase (Zhou et al., 2022), which was recently shown to also turn positive during early MCC differentiation (Basso et al., 2023; Ortiz-Alvarez et al., 2022). Consistent with the p27 observations, while the proportion of pRB+ cells begins to increase in FOXJ1+ WT cells entering A-stage (:: 26%), very few cells are pRB+ in FOXJ1+ *Cena^KO^* cells (:: 3%), a proportion comparable to the proportion observed in WT FOXJ1+ cells at the centrosome stage (:: 4%) (**Fig. 4E and F**). These results suggest that in absence of CCNO, differentiating MCC progenitors do not enter the cell cycle variant and remain in G0/G1-like stage.

Together, these results show that *Cena^KO^* cells committed to becoming MCC stop their differentiation before p27 degradation and RB1 phosphorylation, and suggest that CCNO is necessary to enter the S-like phase of the MCC cell cycle variant.

### Absence of CCNO blocks centriole amplification at the onset of centriole biogenesis

To better characterize the differentiation arrest in *Cena^KO^* cells with regards to centriole biogenesis, we analysed the expression of genes coding for (i) DEUP1, the main component of deuterosomes involved in the scaffolding of centriole biogenesis, (ii) PLK4, the master regulatory kinase involved in centriole assembly, and (iii) SAS6, the cartwheel protein necessary for the onset of centriole assembly. Temporal gene expression analysis in the scRNAseq data show that *Deup1* is successfully activated in *Cena^KO^* cells at the primordial stage. *Plk4* is also activated in mutant cells, although slightly downregulated (**Fig. 3D**), and *Sass6*, predominantly expressed after the primordial stage, remains unexpressed in *Cena^KO^* cells (**Fig. 5A**). These data suggest that, at least at the transcriptional level, *Cena^KO^* cells can prepare for centriole ampliication by expressing *Deup1* and *Plk4* early in the primordial, G0/G1-like phase. The arrest before the expression of *Sass6* suggests that they may not be able to progress further in the centriole biogenesis program.

**Figure 5.**
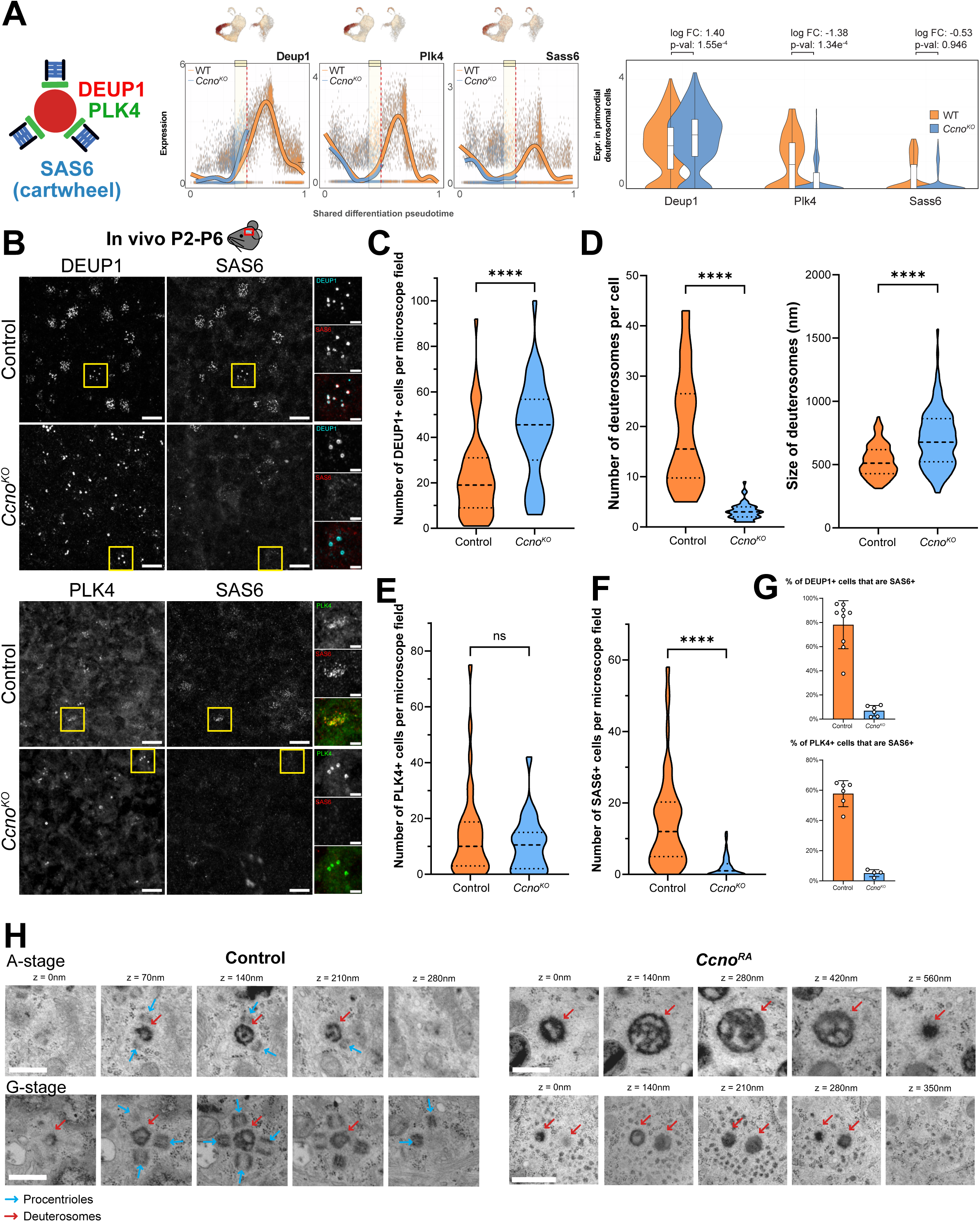
Absence of CCNO blocks centriole amplification at the onset of centriole biogenesis. **(A)** Expression levels of critical centriolar assembly proteins involved in centriole amplification during MCC differentiation. *Deup1* is expressed at similar levels in WT and *Cena^KO^* primordial deuterosomal cells, *Plk4* is slightly downregulated and *Sass6* is poorly expressed in the *Cena^KO^*. **(B)** Tmmunostaining of wholemount lateral ventricles of mice pups aged P2 to P6 for DEUP1, PLK4 and SAS6. *Cena^KO^* cells can express DEUP1 and PLK4 but not SAS6. DEUP1+ and PLK4+ structures are round-shaped suggesting that they are, or they decorate, deuterosomes respectively. Deuterosomes of the *Cena^KO^* are fewer per cell and are enlarged. Scale bar 10µm for wide field images and 2,5µm for zoom-ins. **(C)** Quantification of the number of DEUP1+ cells in microscope field in Control and *Cena^KO^*, showing that *Cena^KO^* have more cells that express DEUP1 compared to the control. Tmages from nine control mice and six *Cena^KO^* mice were quantified, with six images per immunostained tissue, Control: 54 values *Cena^KO^*: 36 values. P-values derived from two-tailed Mann-Whitney U-test, ****p-value<0.0001. **(D)** Quantification of the number of deuterosomes per cell (left) and size of deuterosomes (right) between control and *Cena^KO^*, showing that *Cena^KO^* deuterosomes are less numerous per cell and bigger. Deuterosomes from 3 control animals and 3 *Cena^KO^* animals were counted and measured. Number of deuterosomes: Control: 30 values *Cena^KO^*: 52 values. Size of deuterosomes: Control: 144 values *Cena^KO^*: 128 values. P-values derived from two-tailed Mann-Whitney U-test, ****p-value<0.0001. **(E)** Quantification of the number of PLK4+ cells per microscope field in Control and *Cena^KO^*, showing a similar number of cells expressing PLK4 between control and *Cena^KO^*. Tmages from 6 control mice and 4 *Cena^KO^* mice were quantified, with six images per immunostained tissue, Control: 36 values *Cena^KO^*: 24 values. P-values derived from two-tailed Mann-Whitney U-test, ns: not significant. **(F)** Quantification of the number of SAS6+ cells per microscope field in Control and *Cena^KO^*, showing that *Cena^KO^* fail to express SAS6. Tmages from nine control mice and six *Cena^KO^* mice were quantified, with 6 images per immunostained tissue, Control: 90 values *Cena^KO^*: 60 values. P-values derived from two-tailed Mann-Whitney U-test, ****p-value<0.0001. **(G)** Quantification of the proportion of DEUP1+ cells and PLK4+ cells that can express SAS6 in both WT and *Cena^KO^*. One point represents an animal. **(H)** Electron microscopy images of control and *Cena^RA^* cells in-vitro at DTV5. WT cells show deuterosomes decorated with either A- or G-stage procentrioles. *Cena^RA^* cells show enlarged and empty deuterosomes with no procentrioles. Deuterosomes are indicated by red arrows. Blue arrows indicate procentrioles in the WT. Scale bar 0,5µm.

To confirm whether and how the absence of CCNO blocks centriole biogenesis, we immunostained whole lateral ventricles and cultured ependymal cells for early centriole biogenesis molecular players. *Cena^KO^* cells can express DEUP1 and form deuterosomes *in viva* and *in vitra* (**Fig. 5B-C, Fig. S6A-B**). DEUP1+ cells are even more numerous in *Cena^KO^* cells but form fewer and bigger deuterosomes (**Fig. 5B and D**) suggesting they are blocked during centriole ampliication. Consistent with scRNAseq data, *Cena^KO^*cells also express PLK4 that can take the form of doughnuts suggesting that PLK4 is correctly recruited by deuterosomes (**Fig. 5B and E, Fig. S6A and C**). Of note, fewer *Cena^KO^* cells express PLK4 than DEUP1, which is not the case in the WT cells, suggesting that some cells fail to express, recruit or maintain PLK4 (**Fig. 5C and E**). While the early onset of amplification seems to proceed, nearly no SAS6+ cells are observed *in vitra* or *in viva* in *Cena^KO^* compared to the WT (**Fig. 5B and F, Fig. S6A and D**). No SAS6 is colocalized with DEUP1+ or PLK4+ structures (**Fig. 5B and G, Fig. S6A and E**). These observations are also applicable to *Cena^RA^* cultured cells (**Fig. S6F**). Consistently with the apparent absence of SAS6+ procentrioles, both *Cena^KO^* and *Cena^RA^* cells lack mature centrioles (**Fig. S6G**). Tnterestingly, while we previously showed that overexpression of *Meidas* in *Cena^KO^* cells fails to restore multiciliation, looking at the progression of centriole amplification in this condition revealed that *Meidas* overexpression restores the formation of A-stage SAS6+ procentrioles in a small subset of cells (**Fig. S7**) suggesting that a positive feedback mechanism exists between *Meidas* and *Cena*. Such feedback mechanism could be based on the physical proximity of *Cena* and *Meidas* genetic loci, whose genomic collinearity is conserved across teleosteans (**Fig StA**). However, the procentrioles formed upon *Meidas* overexpression never reach maturity, explaining the absence of multiciliation rescue (**Fig. S5, Fig. S7**).

To test the presence of procentrioles at the ultrastructural level, we used transmission electron microscopy (TEM) on cultured cells of both WT and *Cena^RA^* undergoing MCC differentiation. While WT cells undergoing differentiation show deuterosomes decorated with A- or G-stage procentrioles, no procentrioles are detected on deuterosomes in *Cena^RA^* differentiating cells (**Fig. 5H, Fig. S8**). Tn addition, and consistent with immunostaining data, deuterosomes are fewer, larger and more dense in *Cena^RA^* cells than in WT cells (**Fig. 5D and H, Fig S7F, Fig. S8**).

Taken together, these results shows that mouse brain cells lacking CCNO cannot form the future basal bodies necessary for multicilia formation because of a block in centriole ampliication at the very early onset of centriole assembly.

### Human airway MCC phenotype in the absence of CCNO is similar to mouse brain MCC

*CCNO* was the irst gene found to be mutated (Wallmeier et al., 2014) in a human condition heterogeneously named « acilia syndrome », « ciliary aplasia » or « oligocilia phenotype », and grouping human patients with seemingly bald respiratory epithelia and presenting symptoms similar to Primary Cilia Dyskinesia (PCD). Although patients with RGMC present a higher rate of hydrocephalus compared to the general population, the most debilitating phenotype remains chronic airway infections. The cell cycle variant identified in mouse brain cells in the companion paper seems to be conserved in mouse and human respiratory cells (Serizay et al., 2024). To characterize the role of CCNO in human airway epithelial cells and test whether it is more akin to mouse brain or airways, we generated a hESC H9 line with a 14 bp homozygous deletion in the *CCNO* gene (hereafter referred to as *CCNO*^-/-^) which produces a truncated protein product (**Fig. S9**). Mutation of the *CCNO* gene did not appear to affect the pluripotency or genetic stability of H9 cells (**Fig. S10**). WT and *CCNO*^-/-^ H9 cells were differentiated into airway epithelial cells following a directed differentiation protocol adapted from Hawkins et al., 2021 and Soh et al., 2012. No significant differences were observed in the ability of WT and *CCNO^-1-^* H9 cells to differentiate into lung progenitors and airway basal cells (**Fig. S11**). We then immunostained WT and *CCNO*^-/-^ hESCs-derived airway epithelial cells cultured under air-liquid interface (ALT) conditions for MCC differentiation (FOXJ1, RFX2, RFX3), procentriole (SAS6), and cilia (GT335) markers (**Fig. 6A-C-E, Fig. S12**). This reveals that, while WT and *CCNO*^-/-^ cells can both acquire MCC fate as shown by FOXJ1, RFX2 and RFX3 positivity, only WT cells can produce SAS6+ procentrioles and grow multicilia (**Fig. 6B-D-F, Fig S12**), suggesting that, in absence of CCNO, human airway epithelial cells encounter a block in their differentiation at the early onset of centriole biogenesis, as also found in the mouse brain.

**Figure 6.**
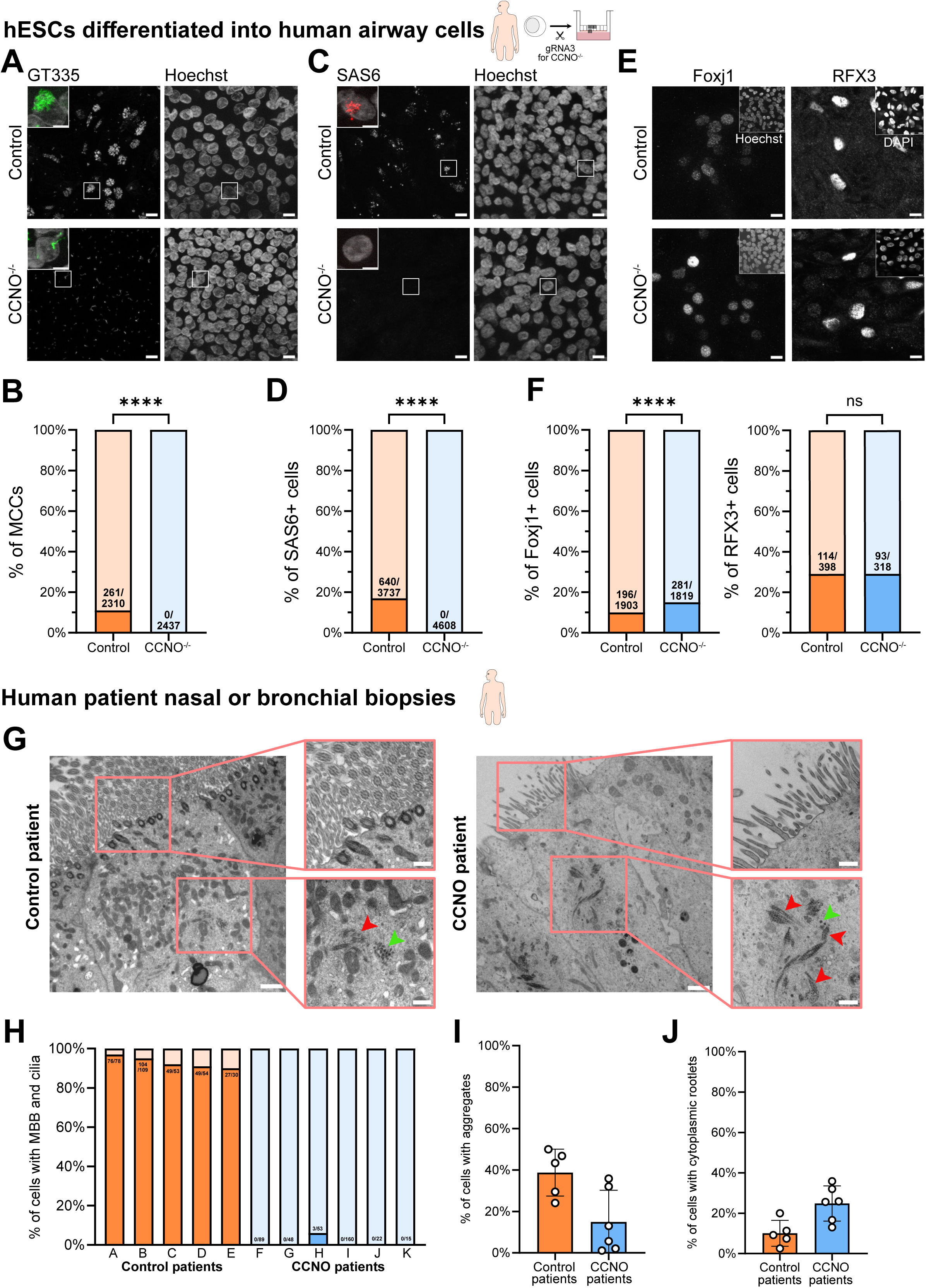
Human airway phenotype in the absence of CCNO is similar to mouse brain MCC. **(A, B)** hESC differentiated into human epithelial airway cells expressing non-functional CCNO (*CCNO*^-/-^) cannot form cilia, marked by GT335. Scale bar 10µm for large-field images and 5µm for zoom-ins. P-values derived from two-sided Chi-square test (two-proportion z-test), ****p-value<0.0001. **(C, D)** hESC differentiated into human epithelial airway cells expressing non-functional CCNO (*CCNO*^-/-^) cannot form SAS6+ procentrioles. Scale bar 10µm for large-field images and 5µm for zoom-ins. P-values derived from two-sided Chi-square test (two-proportion z-test), ****p-value<0.0001. **(E, F)** hESC differentiated into human epithelial airway cells expressing non-functional CCNO (*CCNO*^-/-^) can turn on multiciliogenesis program by expressing FOXJ1 and RFX3. Scale bar 10µm. P-values derived from two-sided Chi-square test (two-proportion z-test), ****p-value<0.0001. **(G)** Airway epithelium of control and CCNO patients in TEM. Scale bar 1µm for large-field images and 0,5µm for zoom-ins. **(H)** Quantification shows that nearly all cells with microvilli show basal bodies (MBB) and cilia in control patients, while nearly no MBB and cilia are observed in the same cells in CCNO patients, as shown in top zoom-in pictures in **(G)**. **(I, J)** Cells with microvilli in human CCNO patients show the presence of centriolar satellites (green arrowheads) and cytoplasmic rootlets (red arrowheads) as indicated in the bottom zoom-in pictures in **(G)**, suggesting that they are halted in the process of centriole ampliication.

To confirm this finding, we analysed nasal brushing biopsies from 15 different patients with mutations in the *CCNO* gene by transmission electron microscopy (**Table S3)**. Previous studies on respiratory MCC from patients with *CCNO* mutations mainly reported cilia scarcity and difficulties in finding basal bodies by electron microscopy (Amirav et al., 2016; Casey et al., 2015; Henriques et al., 2021; Wallmeier et al., 2014). However, the absence of quantification has left unclear whether the phenotype is severe like in the *Cena* mutant mouse brain, or milder, like in its trachea. To identify differentiated cells in patients, we focused our analysis on cells with microvilli, a feature of MCC differentiation. We found that while 94% of cells (305/324) from control patients present multiple basal bodies, nearly no *CCNO* patients’ cell TEM sections (0.1%; 5/420) show more than two centrioles, suggesting that they fail to amplify centrioles and to form basal bodies. Tn line with this, by contrast to control patients’ cells, patients with *CCNO* mutations cells’ do not form multicilia (**Fig. 6G-H, Fig. S13A**). The 3 *CCNO* mutated cells presenting few basal bodies and cilia come from patient H (**Fig. 6H, Fig. S13B**), who carries the frameshift pathogenic variation c.793dup p.(Val265Glyfs*106) in one allele, resulting in a premature Stop codon in the last exon. Transcripts from this allele likely escape nonsense-mediated mRNA decay and may underlie the production of a potentially hypomorphic protein that retains the first cyclin domain. Tnterestingly, in a subset of cells coming from the 15 *CCNO* patients, TEM sections reveal centriolar attributes such as electron-dense aggregates, which are centriolar satellites known to be involved in centriole or cilia formation (Hall et al., 2023; Sorokin, 1968; Zhao et al., 2021) (**Fig. 6G, top zoom-ins and I, Fig. S13A**), and rootlets which constitute the roots of mature centrioles (**Fig. 6G, top zoom-ins and J, Fig. S13A**). This is consistent with scRNAseq of mouse *Cena^KO^* cells showing that mRNA levels of *Pem1* (coding for the core component of centriolar satellites) and *Craee1* (coding for the protein Rootletin, core component of the rootlet) are not changed or are upregulated respectively in deuterosomal primordial cells **(Fig 3D)**. This suggests that, like in mouse brain cells, the *CCNO* mutant human airway MCC prepared for centriole amplification, but were interrupted in the process.

Altogether, these observations suggest that human respiratory MCC need *CCNO* to enter the cell cycle variant and produce centrioles, like mouse brain MCC, and that patients can present mutations leading to the production of a hypomorphic protein that can facilitate the biogenesis of few basal bodies or cilia.

## Discussion

This study shows that Cyclin O controls the entry into the MCC cell cycle variant (described in a companion study by Serizay et al., 2024) and the subsequent centriole amplification required for multiple cilia formation (**Fig. S14**). This work and the associated study argue in favour of CCNO acting as a canonical cyclin, involved in switch-like transitions of a cell cycle variant, rather than an atypical cyclin. First, *Cena* is one of the four cyclins expressed as successive waves along the MCC cycle variant, between *Cend2* and C*enb1/Cena1*, and its expression is temporally correlated with *Cene2* along the canonical cell cycle. Consisently, the expression of *Cena* covers the expression of S-phase and G2/M transition regulators. Second, CCNO depletion leads to the lack of nearly all cells annotated for post-G0/G1 cell cycle-like phases or positive for hallmarks of cell cycle entry and progression (e.g. RB phosphorylation and p27 degradation). Finally, we show that *Cena* is also expressed as a comparable wave, associated with the silencing of *Cene* and with the subsequent expression of *Cenb1* and *Cena1,* during male meiosis. *Cena* is also expressed during female oogenesis, which is involved in meiotic progression (Ma et al., 2013). Altogether, these observations strongly support the role of CCNO as a canonical cyclin, involved in the entry and progression of non-traditional cell cycle variants characterized by the production of centrioles uncoupled from DNA replication.

The earliest steps of centriole biogenesis during the canonical cell cycle occurs at the G1/S transition, concomitant with DNA replication (Nigg and Holland, 2018). Yet, whether and with which temporality the transcription of genes coding for centriole components are regulated needs to be better documented, probably because only two centrioles are produced and transcripts are challenging to detect. During massive amplification of centrioles in MCC, and with single cell transcriptomic resolution, we uncover a transcriptional cascade of core centriole regulators, where centriole scaffolding components *Deup1*, *Cep152* and *PLk4* expression onsets slightly precede early centriolar components such as *Sass6* and *StiL* which precede centriole maturation core regulators such as *Pae5*, *Ninein* and *Cep164*. The expression of some of the genes coding for motile cilia components occurs later. Interestingly, the block in differentiation characterized in the *Cena^KO^* mutant, which occurs before the entry into the S-like phase, follow the onset of expression of centriole scaffolding components and precedes the activation of genes coding for core centriole constituents, from the earliest one, SAS6. This leads to the production of deuterosomes decorated with PLK4, but to the lack of procentrioles, basal bodies and cilia. Supporting the proximity in regulating meiosis and MCC differentiation by CCNO, the block under CCNO depletion is associated with a block in microtubule-organizing centre formation during mouse female meiosis (spindle poles are acentriolar in female meiosis; Ma et al., 2013). Altogether, our data show that the coupling of centriole biogenesis to an S-like phase entry in the MCC cell cycle variant is maintained, as previously suggested by the involvement of Myb in centriole amplification (Tan et al., 2013; Wang et al., 2013) - although DNA does not replicate - and is dependent on CCNO.

In contrast to mouse brain and human airways, in mouse airways and oviducts, CCNO depletion authorizes the formation of around 30% of MCC (Funk et al., 2015; Nt’.nez-Olle et al., 2017, this study). Yet, most of these cells display abnormal ciliation, which suggests that they most likely compensate for the loss of CCNO rather than belonging to a different population, differentiating through a CCNO-independent pathway.

Functional redundancy and compensation for chronic depletion of cyclins or cyclin dependent kinases have been known for a long time (Murray, 2004). Interestingly, this population of cells escaping the most severe phenotype, but later showing defects, reveal a role for CCNO during the late stages of centriole amplification (Funk et al., 2015), consistent with the expression of CCNO during G-stage of centriole biogenesis and G2/M-like stages of the MCC cell cycle variant. Since normal ciliation seems also possible in mutant tissues, it suggests that even the later role of CCNO in centriole maturation and subsequent ciliation can be compensated. *Ccna* is part of a locus containing two other genes involved in multiciliation (*Mcidas* and *Cdc20b*).

Interestingly, their clustering, order and collinearity are conserved among vertebrates (with an exception for zebrafish family), and the temporality of their expression along the pseudotime of differentiation follows their order along the DNA strand, with *Mcidas* and *Ccna* quasi-simultaneously expressed in brain MCC (**Fig. S3C**). This conserved collinearity could play a role in the successive activation of each gene, through the spreading of local chromatin remodelling. Using single-cell resolution transcriptomic measurements, we consistently show that *Ccna* depletion leads to decreased *Mcidas* expression. Nonetheless, it has been demonstrated in other studies that *Mcidas* depletion also blocks *Ccna* expression (Boon et al., 2014; Lu et al., 2019; Wallmeier et al., 2014). We therefore propose a positive feedback loop between *Mcidas* and *Ccna*, whereby *Mcidas* activates *Ccna* expression which in turn activates *Mcidas* expression. Consistent with this hypothesis, *Mcidas* depletion leads to a phenotype comparable to *Ccna* mutant (Boon et al., 2014; Lu et al., 2019; Stubbs et al., 2012), overexpression of *Mcidas* in *Ccna* depleted cells rescues early amplification in a subset of mouse brain cells (this paper), and *Mcidas* depletion is not compensated in mouse airways (Lu et al., 2019). Such positive feedback loop would be reminiscent of G1 cyclins, which increase their transcription to increase cyclin-CDK activity, ensuring commitment to the cell cycle and the activation of the entire G1/S transcriptional network (Bertoli et al., 2013).

## Supporting information

Supplemental informations

## Acknowledgments

We thank all the members of the Spassky lab who contributed to the elaboration of this research work. We also thank the TBENS administrative team and imaging platform for their support and the TBENS Animal Facility for animal care. We thank Kevin Lebrigand and Virginie Magnone for fruitful discussions on single-cell RNA sequencing. We thank Xavier Morin and Stavros Taraviras for sharing plasmids.

This work is supported by funding to A.M. from the ANR (ANR-19-CE13-0027) and Q-life program. The Spassky laboratory is also funded by TNSERM, the CNRS, the Ecole Normale Superieure (ENS), the ANR (ANR-20-CE45-0019, ANR-21-CE16-0016, ANR-22-CE16-0011), the European Research Council (ERC grant agreement 647466) and the Fondation pour la Recherche Medicale (FRM, EQU202103012767). S.R. lab is supported by the Agency for Science, Technology and Research (A*STAR) and a National Medical Research Council (NMRC) of Singapore grant (OFTRG19nov-0037). R.K. is supported by the Institut Pasteur, CNRS, and the European Research Council (ERC grant agreement 771813). J.S. is funded by the Association pour la Recherche sur le Cancer; M.K.D. is funded by the FRM; A.-R.B. is funded by La Ligue Contre Le Cancer; C.T.J. was supported by an A*STAR Research Attachment Program (ARAP) fellowship; S.J.A. is supported by the German Research Foundation (DFG) through a Heisenberg Professorship (AR 732/3-1), project P7 of SFB 1453 (project TD 431984000), and Germany’s Excellence Strategy (CTBSS - EXC-2189 - Project TD 390939984).

This work was performed with supports from the National Tnfrastructure France Genomique (Commissariat aux Grands Tnvestissements, ANR-10-TNBS-09-03, ANR-10-TNBS-09-02), the 3TA Cote d’Azur (ANR-19-P3TA-0002), European Union’s H2020 Research and Tnnovation Program under grant agreement no. 874656 (discovATR), Conseil departemental 06 (2016-294DGADSH-CV).

## Author contributions

Conceptualization: M.K.D., J.S., A.M.; Mouse mutant generation: S.J.A., G.G.G.; Methodology/mouse: M.K.D., J.S., A.-R. B., R.B., M.F., N.D., L.-E.Z.; Single cell formal analysis: J.S.; Methodology/Human patients: M.L., C.F., E.E., D.C.B., H.O.; Methodology/hES-derived MCC: K.J.G., H.L., E.K.T., C.T.J., C.D.B., N.R.D., M.K.D., S.R.; Tnvestigation: M.K.D., J.S., L.-E.Z., P.B., G.G.G., M.L., K.J.G., H.L., E.E., H.O., D.C.B., S.J.A., R.K., S.R., N.S., A.M.; Supervision: A.M.; Writing - original draft: M.K.D., J.S., A.M and edited by all co-authors.

## Data and material availability

All raw and processed sequencing data generated in this study have been submitted to the NCBT Gene Expression Omnibus (GEO; https://www.ncbi.nlm.nih.gov/geo/) and will be publicly released upon publication.

