## Supplemental informations for "Cyclin O controls entry into the cell-cycle variant required for multiciliated cell differentiation"

This file contains:

Supplementary Figures S1 to S14

Supplementary tables S1 to S3

Material and methods

References for methods

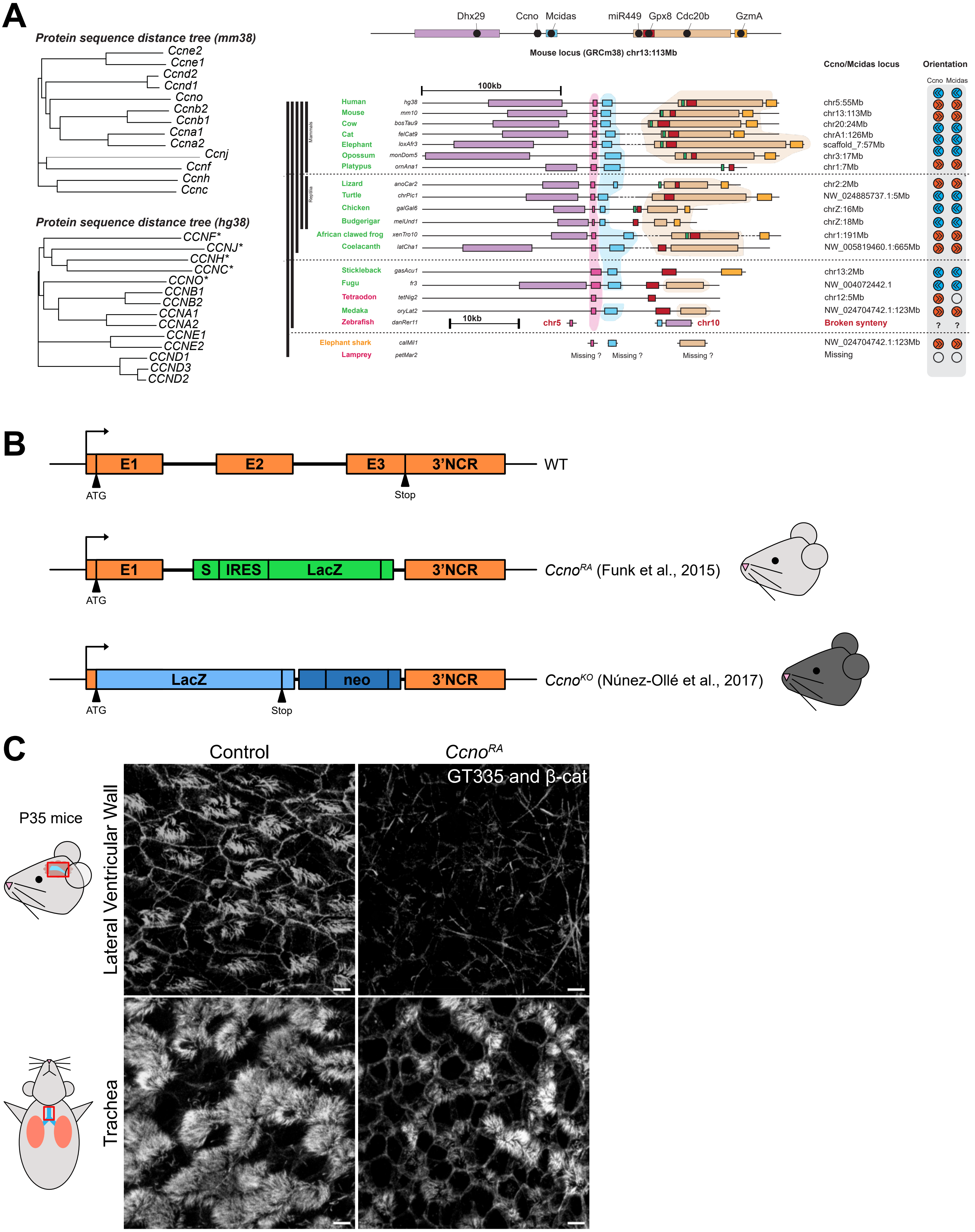

A
Protein sequence distance tree (mm38)
Ccne2
Ccne1
Ccnd2
Ccnd1
Ccno
Ccnb2
Ccnb1
Ccna1
Ccna2
Ccnj
Ccnf
Ccnh
Ccnc
Protein sequence distance tree (hg38)
CCNF*
CCNJ*
CCNH*
CCNC*
CCNO*
CCNB1
CCNB2
CCNA1
CCNA2
CCNE1
CCNE2
CCND1
CCND3
CCND2
Dhx29
Ccno
Mcidas
miR449
Gpx8
Cdc20b
GzmA
Mouse locus (GRCm38) chr13:113Mb
100kb
Ccno/Mcidas locus
Orientation
Ccno
Mcidas
Human hg38 chr5:55Mb
Mouse mm10 chr13:113Mb
Cow bosTau9 chr20:24Mb
Cat felCat9 chrA1:126Mb
Elephant loxAfr3 scaffold_7:57Mb
Opossum monDom5 chr3:17Mb
Platypus ornAna1 chr1:7Mb
Mammals
Lizard anoCar2 chr2:2Mb
Turtle chrPic1 NW_024885737.1:5Mb
Chicken galGal6 chrZ:16Mb
Budgerigar melUnd1 chrZ:18Mb
Reptilia
African clawed frog xenTro10 chr1:191Mb
Coelacanth latCha1 NW_005819460.1:665Mb
Stickleback gasAcu1 chr13:2Mb
Fugu fr3 NW_004072442.1
Tetraodon tetNig2 chr12:5Mb
Medaka oryLat2 NW_024704742.1:123Mb
Zebrafish danRer11 chr5 chr10 Broken synteny
10kb
?
Elephant shark calMil1 NW_024704742.1:123Mb
Lamprey petMar2 Missing
Missing ?
Missing ?
Missing ?
B
E1
E2
E3
3'NCR
WT
ATG
Stop
S
IRES
LacZ
Ccno^RA (Funk et al., 2015)
neo
Ccno^KO (Núnez-Ollé et al., 2017)
C
Control
Ccno^RA
GT335 and β-cat
P35 mice
Lateral Ventricular Wall
Trachea


Figure S1

**Figure S1 – Differential phenotype between brain and trachea exists in both *Ccno*$^{KO}$ and *Ccno*$^{RA}$ mutants.**

**(A)** Phylogeny of CCNO protein in both mouse and human and conservation of CCNO gene locus among various species.

**(B)** *Ccno* gene alleles in WT, *Ccno*$^{KO}$ and *Ccno*$^{RA}$ mouse lines. *Ccno*$^{RA}$ has two exons removed and *Ccno*$^{KO}$ has all exons removed.

**(C)** Immunostaining of LVW and trachea dissected from a single *Ccno*$^{RA}$ or WT individual at P35 for cilia (GT335) and β-cat, showing a complete lack of MCC in the LVW and a decrease of MCC in the trachea, similar to what is observed in *Ccno*$^{KO}$. Scale bar 5µm.

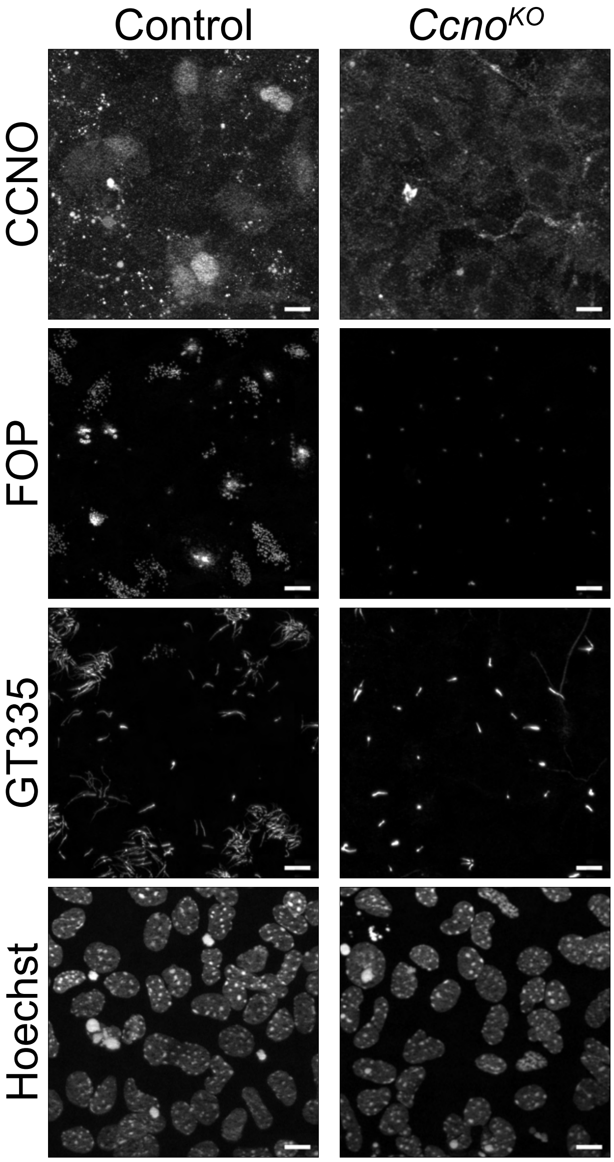


**Figure S2**

**Figure S2 – Validation of CCNO antibody in *Ccno*$^{KO}$ cultured ependymal cells at DIV2.** Scale bar 10µm.

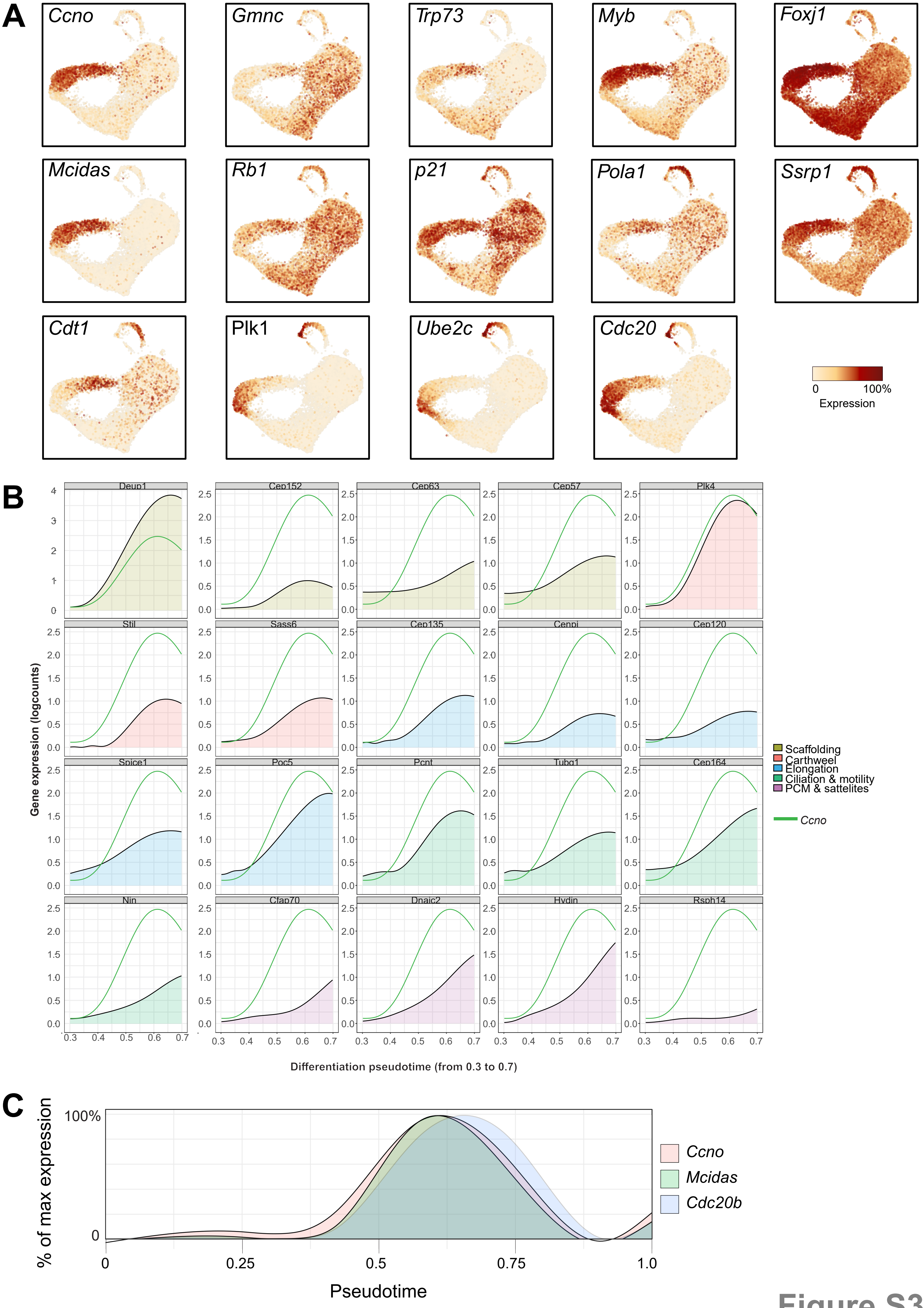
A
Ccno
Gmnc
Trp73
Myb
Foxj1
Mcidas
Rb1
p21
Pola1
Ssrp1
Cdt1
Plk1
Ube2c
Cdc20
0
100%
Expression
B
Deup1
Cep152
Cep63
Cep57
Plk4
Stil
Sass6
Cep135
Cenpj
Cep120
Spice1
Poc5
Pcnt
Tubg1
Cep164
Nin
Cfap70
Dnaic2
Hydin
Rsph14
Gene expression (logcounts)
Differentiation pseudotime (from 0.3 to 0.7)
Scaffolding
Carthweel
Elongation
Ciliation & motility
PCM & sattelites
Ccno
C
% of max expression
100%
0
0.25
0.5
0.75
1.0
Pseudotime
Ccno
Mcidas
Cdc20b

Figure S3

**Figure S3 – Cyclin O is expressed at the onset of MCC differentiation, MCC ell cycle variant and MCC centriole amplification.**

**(A)** UMAP plots of gene expression listed in Fig. 2H-I-J

**(B)** Expression of various genes involved in centriole biogenesis and ciliation compared to *Ccno* expression along differentiation pseudotime.

**(C)** Expression of *Ccno,* Mcidas and *Cdc20b* along differentiation pseudotime.

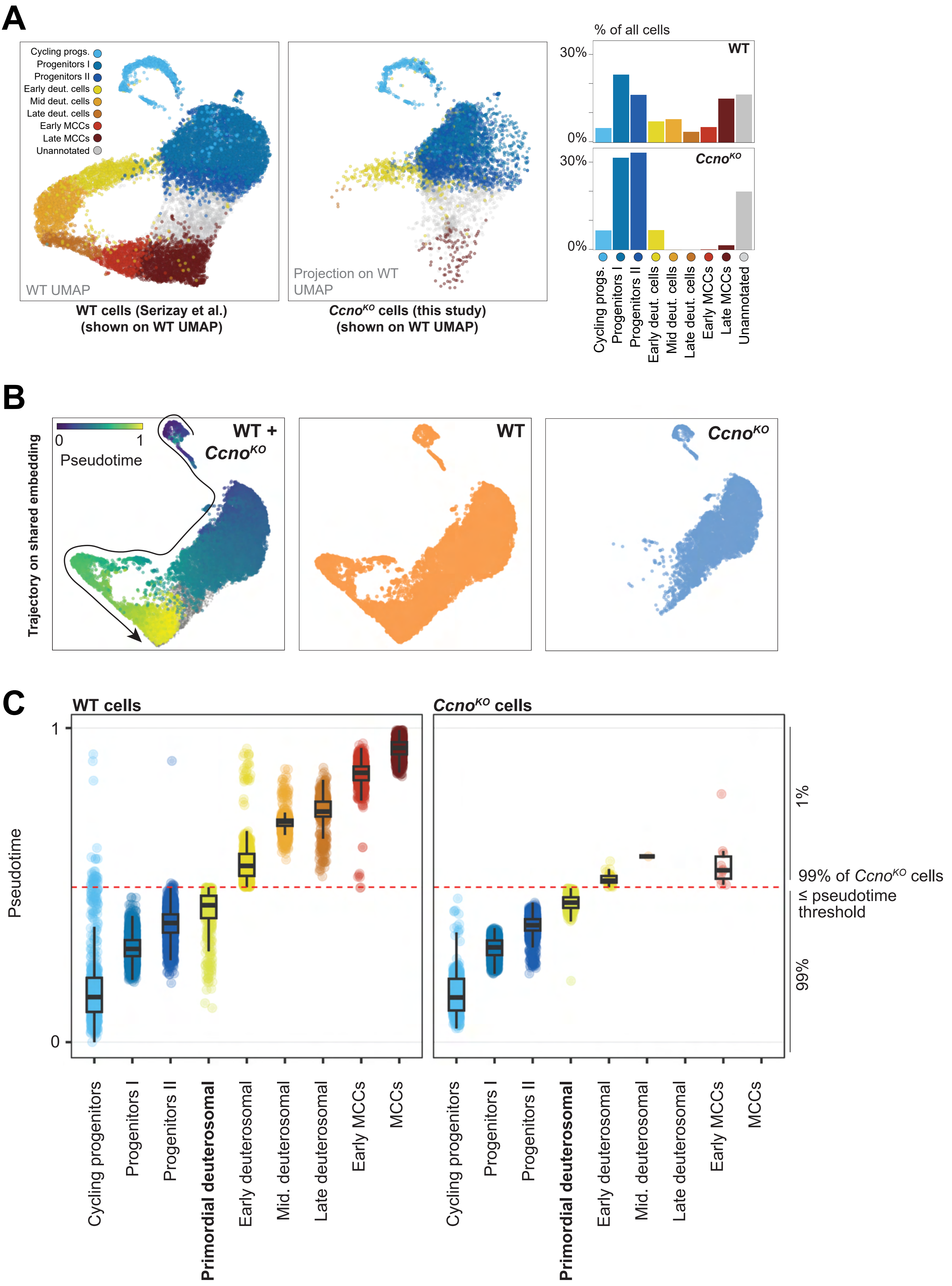
A
Cycling progs.
Progenitors I
Progenitors II
Early deut. cells
Mid deut. cells
Late deut. cells
Early MCCs
Late MCCs
Unannotated
WT UMAP
WT cells (Serizay et al.)
(shown on WT UMAP)
Projection on WT UMAP
Ccno^KO cells (this study)
(shown on WT UMAP)
% of all cells
30%
0%
WT
Ccno^KO
B
Pseudotime
0
1
Trajectory on shared embedding
WT + Ccno^KO
WT
Ccno^KO
C
WT cells
Ccno^KO cells
Pseudotime
1
0
Cycling progenitors
Progenitors I
Progenitors II
Primordial deuterosomal
Early deuterosomal
Mid. deuterosomal
Late deuterosomal
Early MCCs
MCCs
1%
99%
99% of Ccno^KO cells
≤ pseudotime
threshold

Figure S4

**Figure S4 – scRNAseq dataset of *Ccno*$^{KO}$ cells**

**(A)** UMAP projections of WT and *Ccno*$^{KO}$ scRNAseq datasets, with annotations transferred from Serizay et al., 2024.

**(B)** Lineage inference and pseudotime computation of combined WT and *Ccno*$^{KO}$ datasets.

**(C)** Thresholding of pseudotime used to determine the primordial deuterosomal cluster.

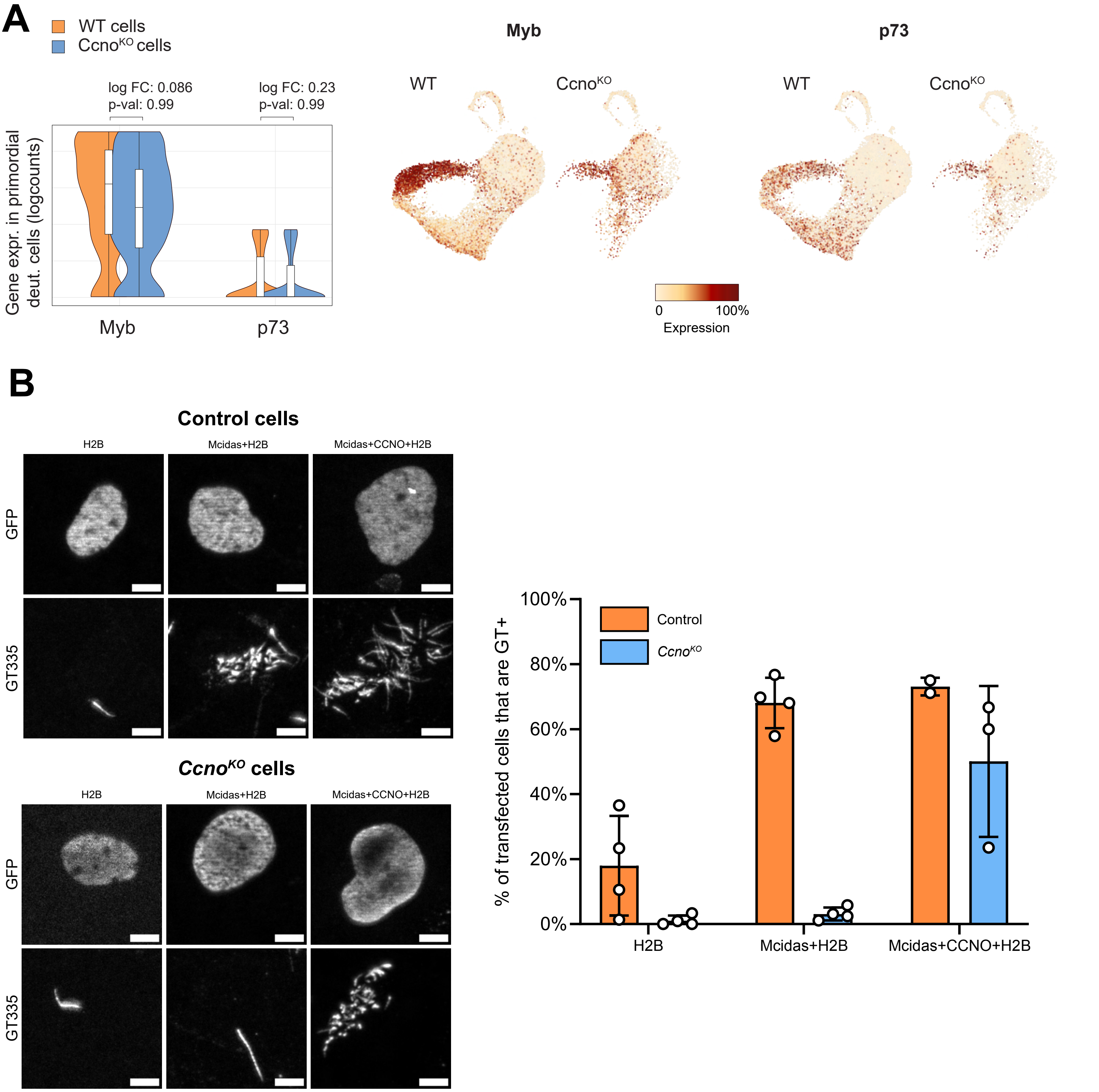

A
WT cells
Ccno^KO cells
log FC: 0.086
p-val: 0.99
log FC: 0.23
p-val: 0.99
Gene expr. in primordial deut. cells (logcounts)
Myb
p73
Myb
p73
WT
CcnoKO
WT
CcnoKO
0
100%
Expression
B
Control cells
H2B
Mcidas+H2B
Mcidas+CCNO+H2B
GFP
GT335
CcnoKO cells
H2B
Mcidas+H2B
Mcidas+CCNO+H2B
GFP
GT335
% of transfected cells that are GT+
100%
80%
60%
40%
20%
0%
Control
CcnoKO
H2B
Mcidas+H2B
Mcidas+CCNO+H2B


**Figure S5**

**Figure S5 – Mcidas overexpression in *Ccno*$^{KO}$ cells fails to rescue multiciliation.**

**(A)** Expression of MCC differentiation factors Myb and p73 in primordial deuterosomal cells from WT and *Ccno*$^{KO}$ samples. Both genes are expressed at comparable levels in WT and *Ccno*$^{KO}$.

**(B)** Transfection of control and *Ccno*$^{KO}$ cells by Mcidas or Mcidas+Ccno, followed by GT335 immunostaining at DIV5. Scale bar 5µm. One point represents an experiment.

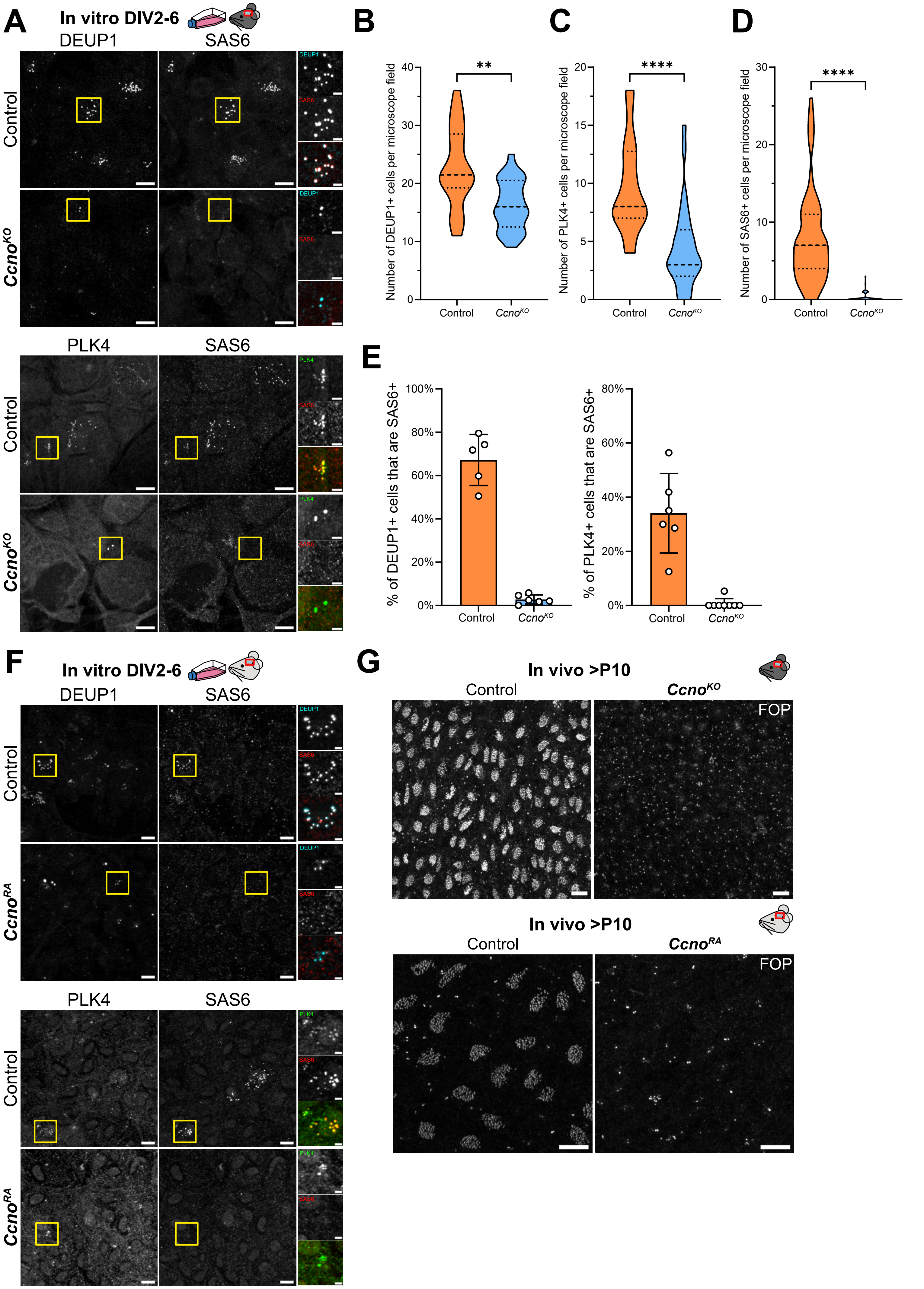

A
In vitro DIV2-6
DEUP1
SAS6
Control
Ccno^KO
PLK4
SAS6
B
Number of DEUP1+ cells per microscope field
**
Control
Ccno^KO
C
Number of PLK4+ cells per microscope field
****
D
Number of SAS6+ cells per microscope field
****
E
% of DEUP1+ cells that are SAS6+
% of PLK4+ cells that are SAS6+
F
In vitro DIV2-6
Ccno^RA
G
In vivo >P10
FOP


Figure S6

**Figure S6 – Absence of CCNO blocks centriole amplification at the onset of centriole biogenesis**

**(A)** Immunostaining of in-vitro cultures at ages between DIV2 and DIV6 for DEUP1, PLK4 and SAS6. $Ccno^{KO}$ cells are able to express DEUP1 and PLK4 but not SAS6. Scale bar 10µm for large field images and 2,5µm for zoom-ins.

**(B)** Quantification of the number of DEUP1+ cells in microscope field in Control and $Ccno^{KO}$ in vitro. Images from 5 control mice and 6 $Ccno^{KO}$ mice were quantified, with 4 images per cultured coverslip, Control: 20 values $Ccno^{KO}$: 25 values. P-values derived from two-tailed Mann-Whitney U-test, **p-value=0.0013.

**(C)** Quantification of the number of PLK4+ cells in microscope field in Control and $Ccno^{KO}$ in vitro. Images from 6 control mice and 8 $Ccno^{KO}$ mice were quantified, with 4 images per cultured coverslip, Control: 24 values $Ccno^{KO}$: 31 values. P-values derived from two-tailed Mann-Whitney U-test, ****p-value<0.0001.

**(D)** Quantification of the number of SAS6+ cells in microscope field in Control and $Ccno^{KO}$ in vitro, showing that $Ccno^{KO}$ fail to express SAS6. Images from 12 control mice and 12 $Ccno^{KO}$ mice were quantified, with 4 to 8 images per cultured coverslip, Control: 68 values $Ccno^{KO}$: 75 values. P-values derived from two-tailed Mann-Whitney U-test, ****p-value<0.0001.

**(E)** Quantification of the proportion of DEUP1+ cells and PLK4+ cells that are able to express SAS6 in both WT and $Ccno^{KO}$ in vitro. One point represents an animal.

**(F)** Immunostaining of in-vitro cultures of $Ccno^{RA}$ at DIV3 for DEUP1, PLK4 and SAS6. $Ccno^{KO}$ cells are able to express DEUP1 and PLK4 but not SAS6. Scale bar 10µm for large field images and 2,5µm for zoom-ins.

**(G)** Immunostaining of in-vivo controls and $Ccno^{KO}$ or $Ccno^{RA}$ lateral ventricles (>P10) for basal bodies, marked by FOP. Scale bar 10µm.

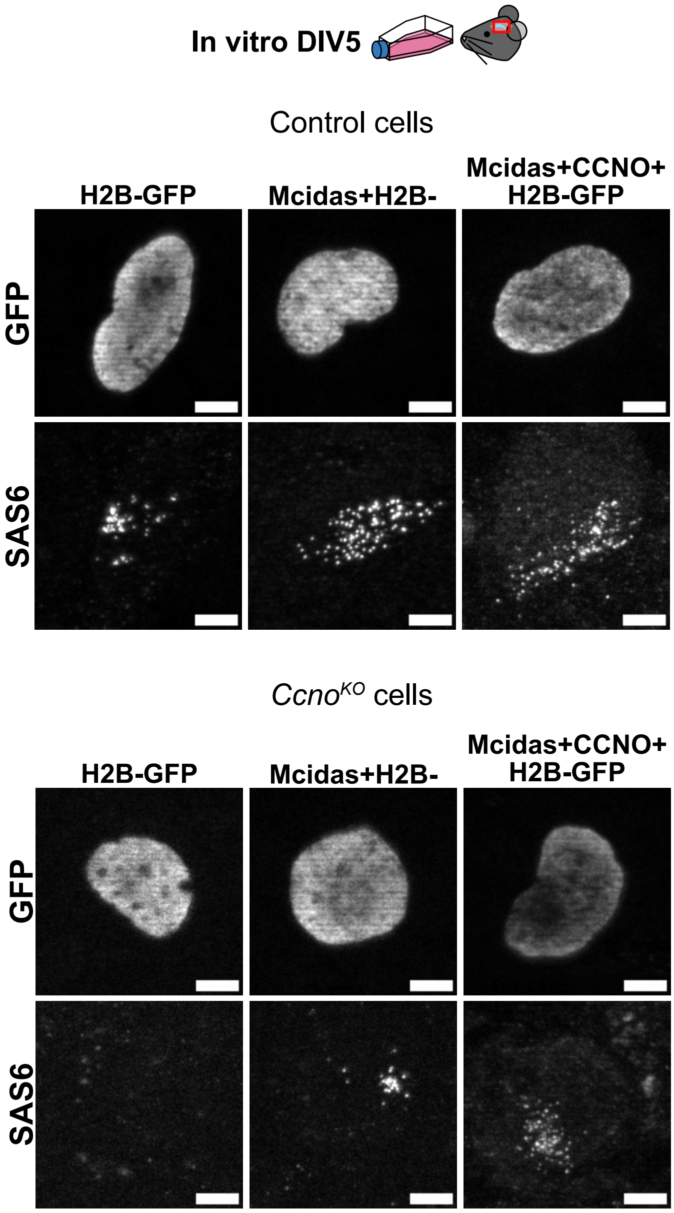

In vitro DIV5
Control cells
H2B-GFP
Mcidas+H2B-
Mcidas+CCNO+
H2B-GFP
GFP
SAS6
*Ccno*KO cells
H2B-GFP
Mcidas+H2B-
Mcidas+CCNO+
H2B-GFP
GFP
SAS6


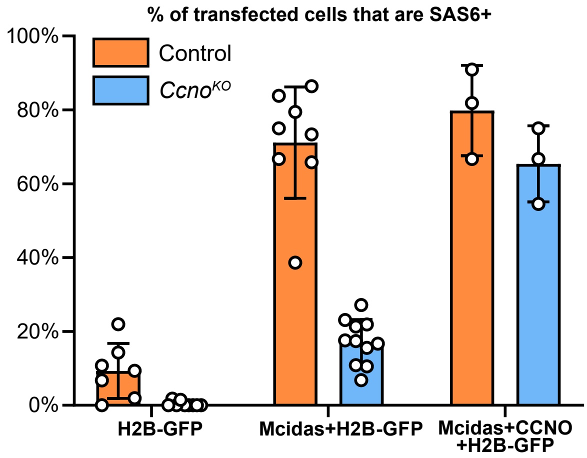

% of transfected cells that are SAS6+
100%
80%
60%
40%
20%
0%
Control
*Ccno*KO
H2B-GFP
Mcidas+H2B-GFP
Mcidas+CCNO
+H2B-GFP


Figure S7

**Figure S7 - Mcidas overexpression in *Ccno*$^{KO}$ partially rescue SAS6 expression.** Controls and *Ccno*$^{KO}$ cells are transfected with plasmid(s) coding for H2B-GFP, H2B-GFP+MCIDAS or H2B-GFP+MCIDAS+CCNO at DIV-1, and immunostained at DIV5 for SAS6 and GFP. Scale bar 5µm. One point represents an experiment.

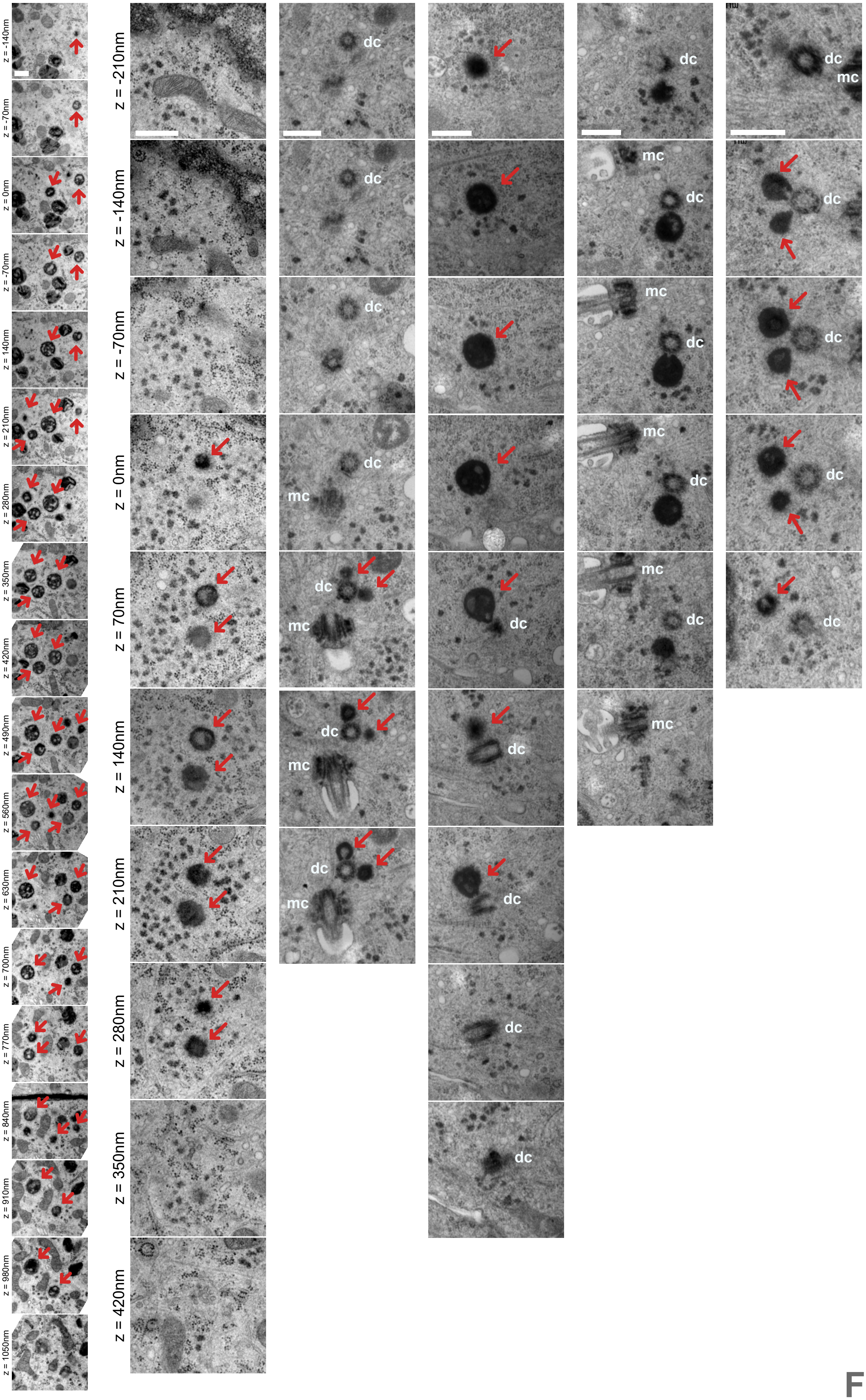
z = -140nm
z = -70nm
z = 0nm
z = -70nm
z = 140nm
z = 210nm
z = 280nm
z = 350nm
z = 420nm
z = 490nm
z = 560nm
z = 630nm
z = 700nm
z = 770nm
z = 840nm
z = 910nm
z = 980nm
z = 1050nm
z = -210nm
z = -140nm
z = -70nm
z = 0nm
z = 70nm
z = 140nm
z = 210nm
z = 280nm
z = 350nm
z = 420nm
dc
mc

Figure S8

**Figure S8 - Electron microscopy images show enlarged and empty deuterosomes with no procentrioles in *Ccno$^{RA}$* cells.** Serial ultra-thin sections of cell from *Ccno$^{RA}$* cells at DIV5 showing empty deuterosomes in the cytoplasm, or connected to the daughter centriole of the centrosome. Deuterosomes are indicated by red arrows. mc: mother centriole of the centrosome, dc: daughter centriole of the centrosome. Scale bar 0,5µm.

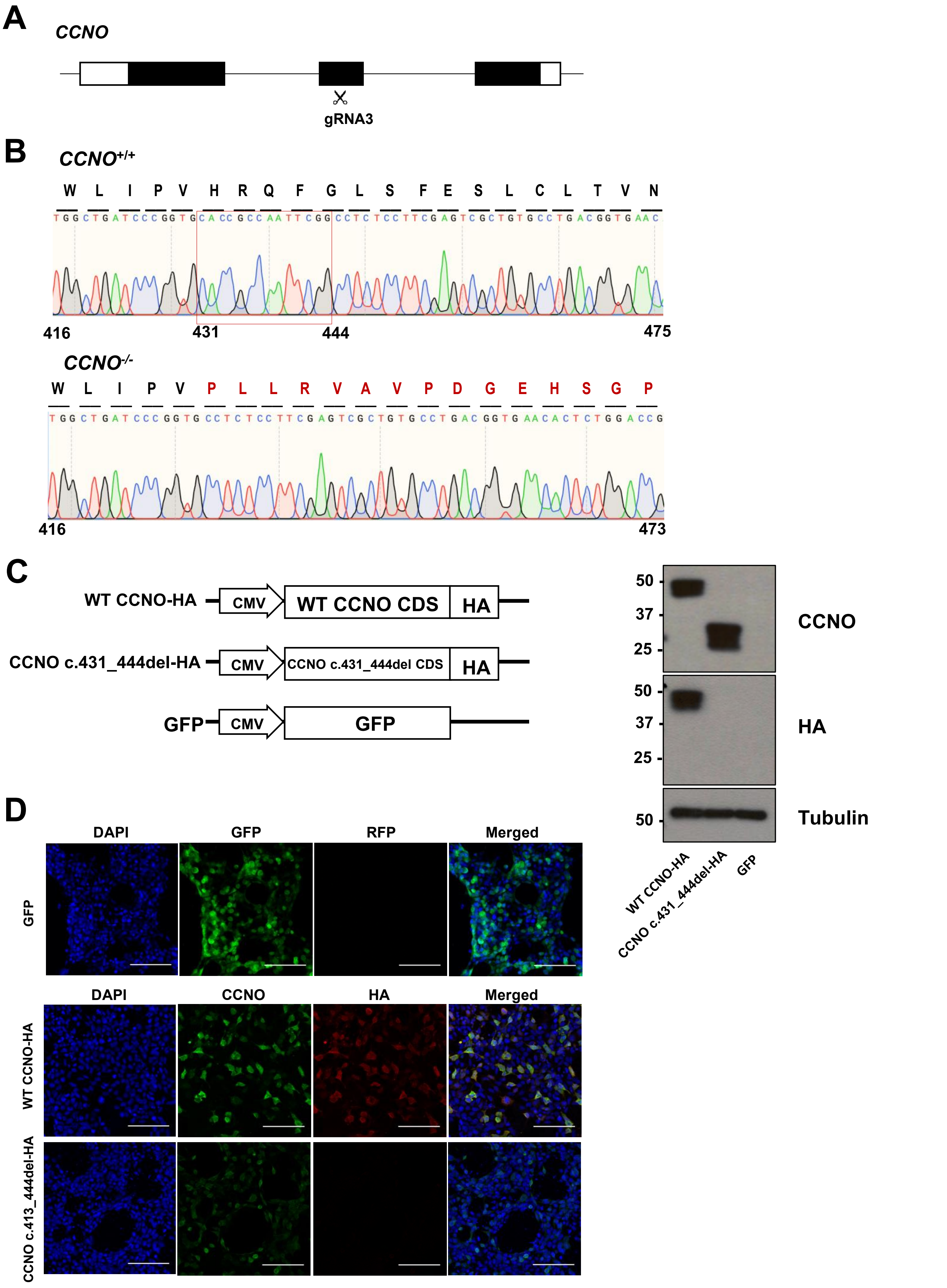

A
CCNO
gRNA3
B
CCNO+/+
W L I P V H R Q F G L S F E S L C L T V N
TGGCTGATCCCGGTGCACCGCCAATTCGGCCTCTCCTTCGAGTCGCTGTGCCTGACGGTGAAC
416
431
444
475
CCNO-/-
W L I P V P L L R V A V P D G E H S G P
TGGCTGATCCCGGTGCCTCTCCTTCGAGTCGCTGTGCCTGACGGTGAACACTCTGGACCG
416
473
C
WT CCNO-HA
CMV
WT CCNO CDS
HA
CCNO c.431_444del-HA
CMV
CCNO c.431_444del CDS
HA
GFP
CMV
GFP
50
37
25
CCNO
50
37
25
HA
50
Tubulin
WT CCNO-HA
CCNO c.431_444del-HA
GFP
D
DAPI
GFP
RFP
Merged
GFP
DAPI
CCNO
HA
Merged
WT CCNO-HA
CCNO c.413_444del-HA


Figure S9

**Figure S9 – Generation of CCNO$^{-/-}$ H9 ESCs.**

**(A)** Schematic of gene editing strategy. gRNA3 was selected which targets exon 2 of the *CCNO* gene.

**(B)** Sanger sequencing of the *CCNO* gene in *CCNO*$^{+/+}$ and *CCNO*$^{-/-}$ cells. Red box indicate the 14 bp sequence deletion found in the *CCNO*$^{-/-}$ cells.

**(C)** Western blotting of CCNO or HA in HEK293T cells overexpressing HA-tagged wild type (WT) and mutant CCNO cDNA sequences. Tubulin was used as a loading control. GFP-overexpressing HEK293T cells was used as a negative control.

**(D)** Representative images of HEK293T cells overexpressing HA-tagged wild type (WT) and mutant CCNO cDNA sequences immunostained for CCNO or HA. GFP-overexpressing HEK293T cells was used as a negative control.

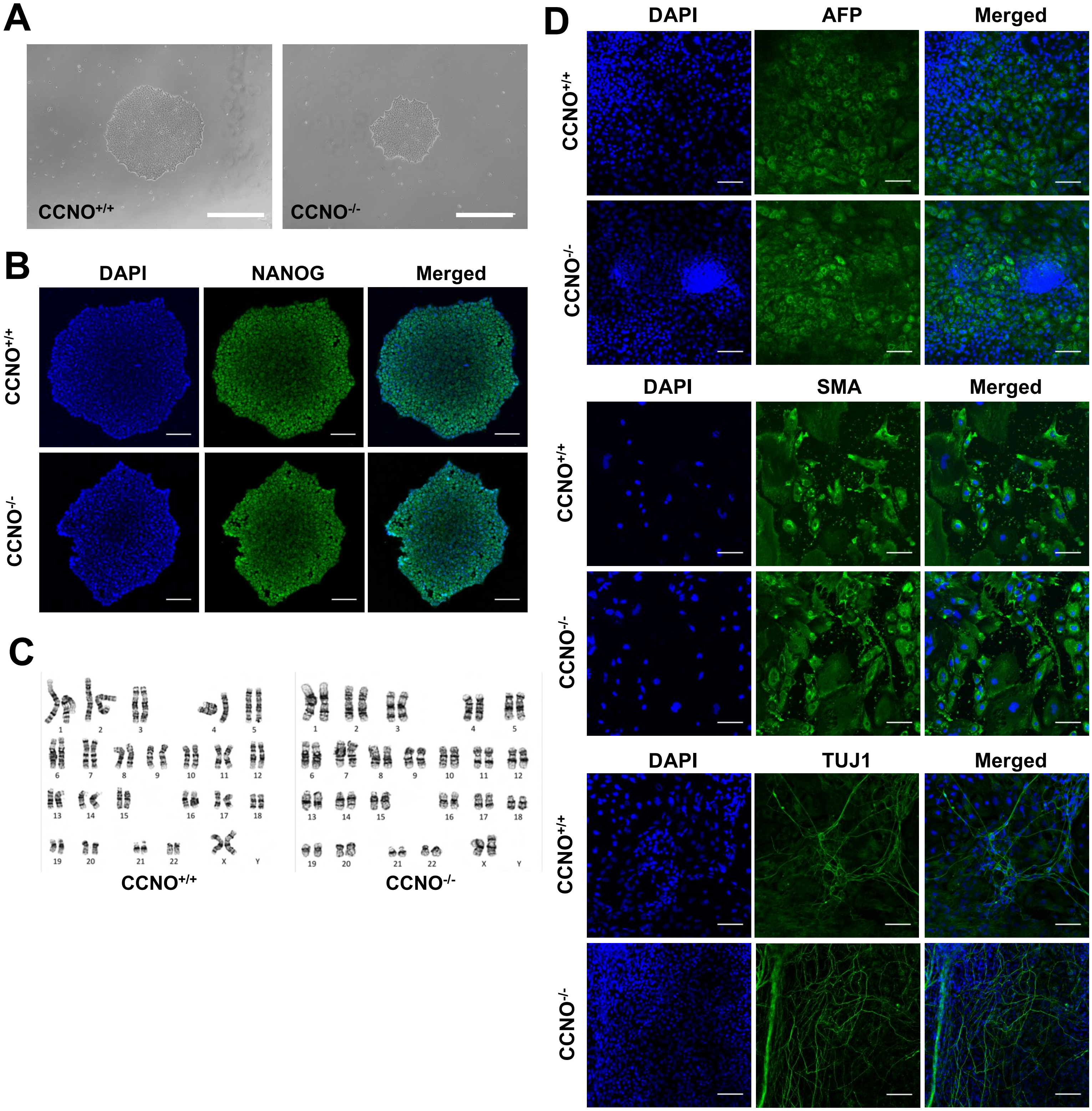
A
CCNO+/+
CCNO-/-
B
DAPI
NANOG
Merged
CCNO+/+
CCNO-/-
C
CCNO+/+
CCNO-/-
D
DAPI
AFP
Merged
CCNO+/+
CCNO-/-
DAPI
SMA
Merged
CCNO+/+
CCNO-/-
DAPI
TUJ1
Merged
CCNO+/+
CCNO-/-

Figure S10

**Figure S10 – CCNO$^{-/-}$ H9 ESCs are pluripotent and retain a normal karyotype.**

**(A)** Brightfield images of representative *CCNO*$^{+/+}$ and *CCNO*$^{-/-}$ ESC colonies.

**(B)** Representative images of *CCNO*$^{+/+}$ and *CCNO*$^{-/-}$ ESC immunostained for pluripotency marker NANOG.

**(C)** Karyotype analysis of *CCNO*$^{+/+}$ and *CCNO*$^{-/-}$ ESCs.

**(D)** Trilineage differentiation of *CCNO*$^{+/+}$ and *CCNO*$^{-/-}$ ESCs followed by immunostaining of markers specific to the 3 germ layers: AFP (endoderm), SMA (mesoderm) and TUJ1 (ectoderm).

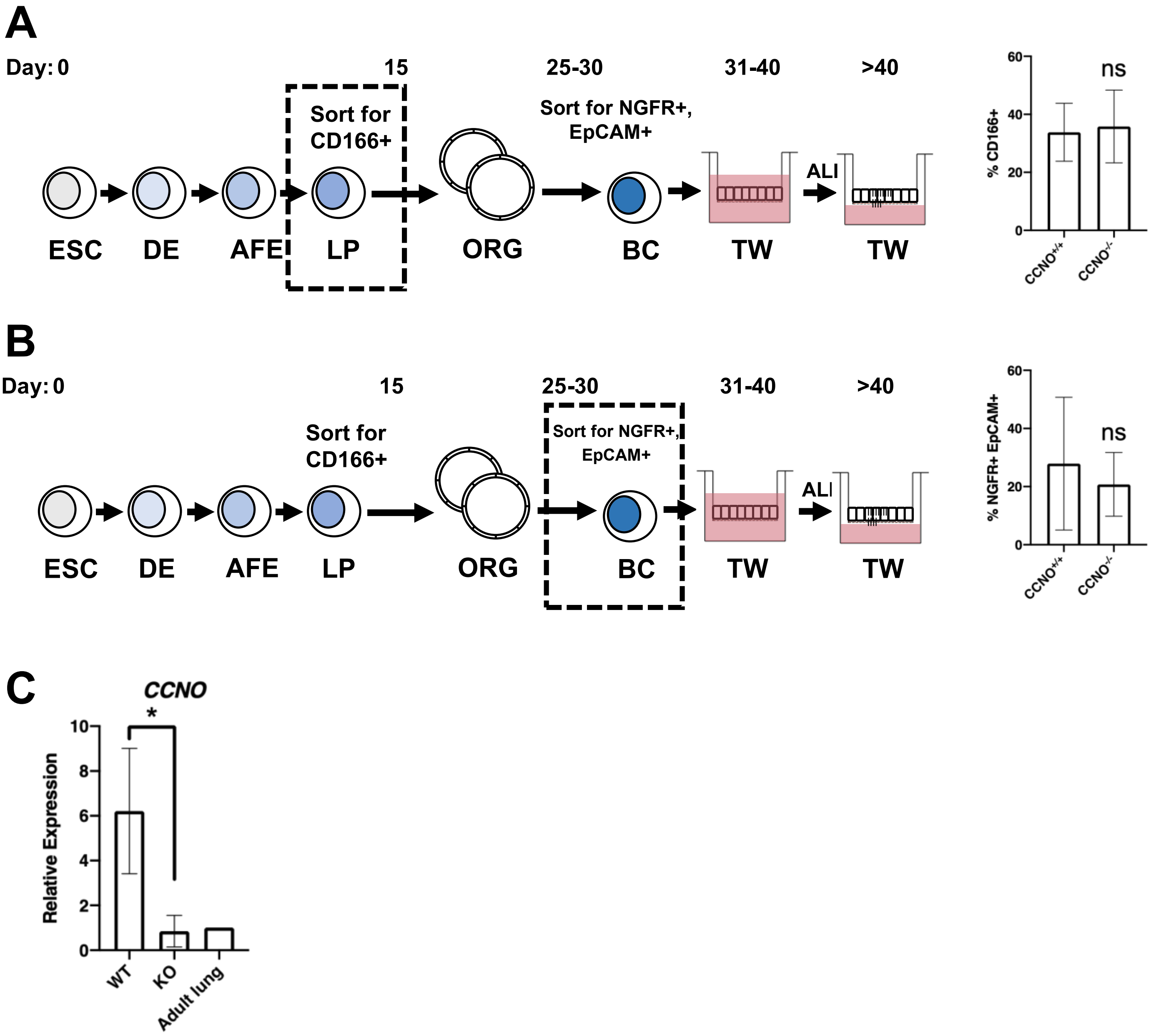
A
Day: 0
15
25-30
31-40
>40
Sort for CD166+
Sort for NGFR+, EpCAM+
ALI
ESC
DE
AFE
LP
ORG
BC
TW
TW
% CD166+
60
40
20
0
ns
CCNO+/+
CCNO-/-
B
Day: 0
15
25-30
31-40
>40
Sort for CD166+
Sort for NGFR+, EpCAM+
ALI
ESC
DE
AFE
LP
ORG
BC
TW
TW
% NGFR+ EpCAM+
60
40
20
0
ns
CCNO+/+
CCNO-/-
C
CCNO
*
Relative Expression
10
8
6
4
2
0
WT
KO
Adult lung

Figure S11

**Figure S11 – Stepwise differentiation of $CCNO^{+/+}$ and $CCNO^{-/-}$ ESCs into airway epithelial cells.**

**(A)** Schematic of differentiation protocol. Black box indicates when cells were sorted and assessed for % CD166 expression in lung progenitors (LP).

**(B)** Schematic of differentiation protocol. Black box indicates when cells were sorted and assessed for % NGFR;EpCAM expression in proximal lung organoids (ORG).

**(C)** QPCR analysis of proximal airway epithelial cell markers. n=3. Expression was normalized to adult lung cDNA. *p <0.05. Student's t test.

**A**

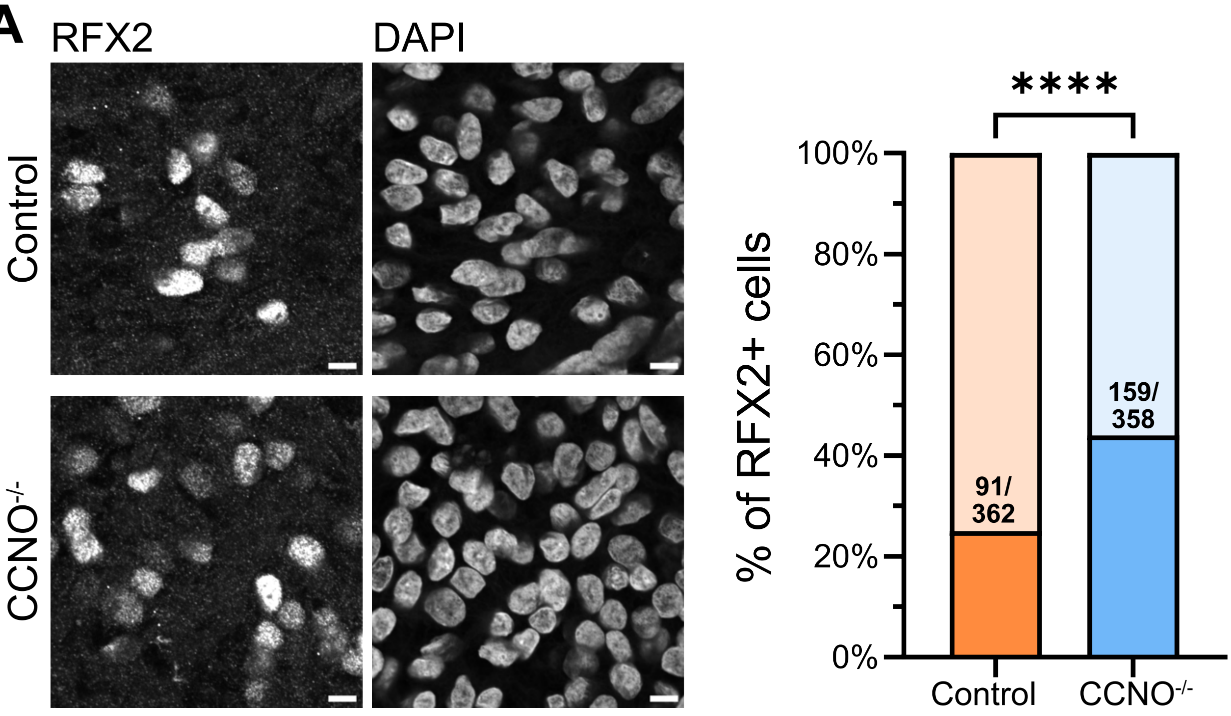
RFX2
DAPI
Control
CCNO-/-
****
100%
80%
60%
40%
20%
0%
% of RFX2+ cells
91/
362
159/
358
Control
CCNO-/-

**Figure S12**

**Figure S12 – RFX2 expression in control vs CCNO$^{-/-}$ ESCs differentiated into airway epithelial cells.** Scale bar 5µm. P-values derived from two-sided Chi-square test (two-proportion z-test), ****p-value<0.0001.

**A**

| CCNO patient | Total cells | Cells with aggregates | Cells with cytoplasmic rootlets | Cells with more than 2 BB/cilia |
|---|---|---|---|---|
| L | 6 | 0 | 0 | 0 |
| M | 3 | 0 | 1 | 2 |
| N | 4 | 0 | 0 | 0 |
| O | 4 | 3 | 0 | 0 |
| P | 1 | 1 | 0 | 0 |
| Q | 7 | 1 | 2 | 0 |
| R | 4 | 2 | 3 | 0 |
| S | 1 | 0 | 0 | 0 |
| T | 3 | 0 | 0 | 0 |

**B**

CCNO patient H

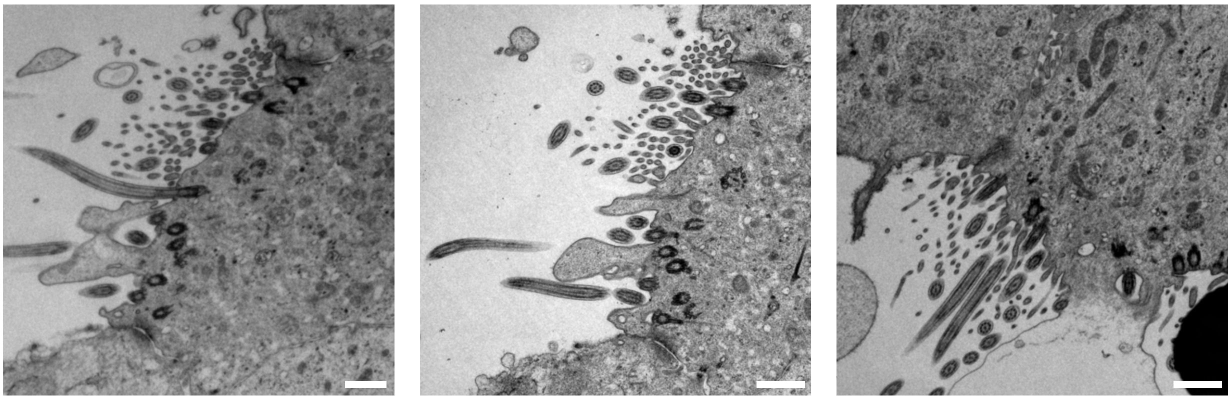

**Figure S13**

**Figure S13 – Transmission electron microscopy images of CCNO human patients**

**(A)** Quantification of numbers of cells that have rootlets, aggregates and more than two basal bodies and/or cilia in CCNO patients listed.

**(B)** TEM images of the 3 cells (out of 53) from CCNO patient H showing multiple basal bodies and/or cilia. Scale bar 1μm.

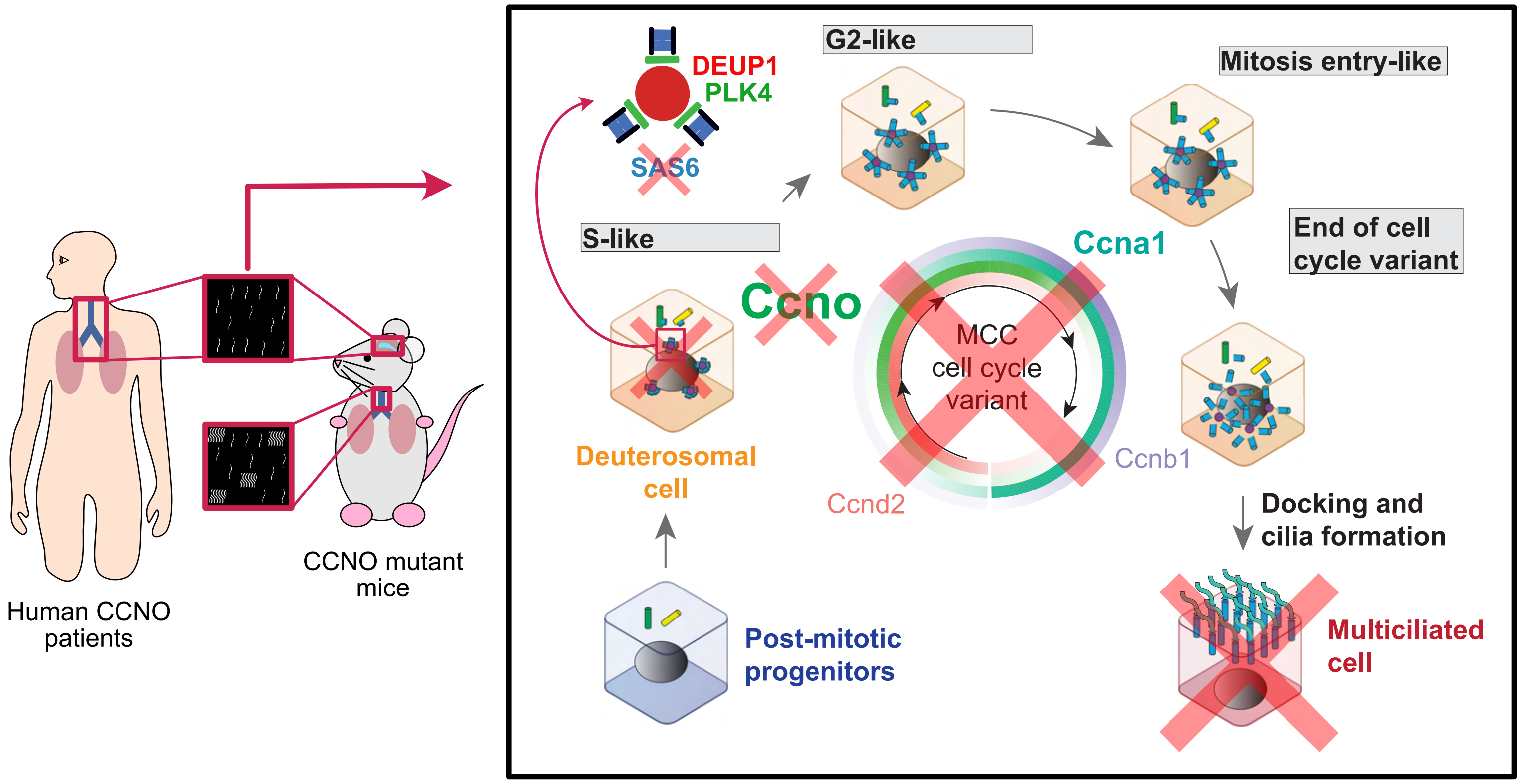
DEUP1
PLK4
SAS6
G2-like
Mitosis entry-like
S-like
Ccno
Ccna1
End of cell cycle variant
MCC cell cycle variant
Deuterosomal cell
Ccnd2
Ccnb1
Docking and cilia formation
Post-mitotic progenitors
Multiciliated cell
Human CCNO patients
CCNO mutant mice

Figure S14

**Figure S14 – Cyclin O is necessary for multiciliated cells to enter their differentiation cell cycle variant and al-lows the massive amplification of centrioles, which serve as basal bodies for cilia nucleation.**

**Table S1.** gRNA and primer sequences

| **gRNA/primer** | **Sequence (5'->3')** | **Description** |
|---|---|---|
| hCCNO sgRNA3 | ACTCGAAGGAGAGGCCGAATTGG | CRISPR sgRNA3 |
| hCCNO F1 | atggtgaccccctgtccca | Genotyping |
| hCCNO R1 | ttatttcgagctcgggggcag | Genotyping |
| EcoRI-Flag-hCCNO | GACGACGATGACAAGgtgaccccctgtcccac | Insertion of FLAG tag at N terminus |
| EcoRI-Flag-hCCNO | CATGGACTACAAGGACGACGATGACAAGgtgacc | Insertion of FLAG tag at N terminus |
| EcoRI-Flag-hCCNO | tcagGAATTCGCCACCATGGACTACAAGGACGAC- | Insertion of FLAG tag at N terminus |
| hCCNO-XbaI R1 | tcagTCTAGAttatttcgagctcgggggcag | Insertion of FLAG tag at N terminus |
| SacI-Ha-XbaI Oligo 1 | cgaaaTACCCATACGATGTTCCAGATTACGCTtaaT | Insertion of HA tag at C terminus |
| SacI-Ha-XbaI Oligo 2 | CTAGAttaAGCGTAATCTGGAACATCGTATGGG- | Insertion of HA tag at C terminus |
| **RT qPCR primer** | **Forward sequence (5'->3')** | **Reverse sequence (5'->3')** |
| ACTB | CTGGAACGGTGAAGGTGACA | AAGGGACTTCCTG- |
| CCNO | TCTACAGACCTTCCGCGACT | TCCAGAGTGTTCACCGTCAG |

**Table S2.**

| Primary antibodies | | | | | |
|---|---|---|---|---|---|
| **Name** | **Species** | **Application** | **Company** | **Catalogue no./reference** | **Dilution** |
| Anti-polyglutamylated tubulin antibody (GT335) | Mouse | IF | Adipogen | AG-20B-0020-C100 | 1:1000 |
| Anti-Acitve-β-Catenin antibody (Clone 8E7) | Mouse | IF | Sigma-Aldrich | 05-665 | 1:500 |
| Anti-CCNO antibody | Rabbit | IF | Homemade | Gabriel Gil-Gómez lab | 1:20 |
| Anti-FOP antibody (FGFR1OP, Clone 2B1) | Mouse | IF | Abnova | H00011116-M01 | 1:1000 |
| Anti-FoxJ1 antibody (Clone 2A5) | Mouse | IF | eBioscience | 14-9965 | 1:700 |
| Anti-p27kip1 | Rabbit | IF | Santa Cruz | sc-528 | 1:200 |
| Anti-Phospho-Rb Ser807/811 antibody (Clone D20B12) | Rabbit | IF | Cell Signaling | CST 9308 | 1:1000 |
| Anti-Deup1 (raised against TKLKQSRHI peptide) | Rabbit | IF | Homemade | Mercey et al., 2019 | 1:5000 |
| Anti-SAS6 antibody (91.390.21) | Mouse | IF | Santa Cruz | sc-81431 | 1:700 |
| Anti-Plk4 antibody coupled to AF555 | Rabbit | IF | Homemade | LoMastro et al., 2022 | 1:1000 |
| ALCAM Phycoerythrin monoclonal antibody (Clone 105902) | Mouse | Flow cytom-etry | R&D Systems | FAB6561P | 1:200 |
| APC anti-human CD271 (NGFR) antibody | Mouse | Flow cytom-etry | Biolegend | 345108 | 1:200 |
| PE anti-human CD326 (EpCAM) antibody | Mouse | Flow cytom-etry | Biolegend | 324206 | 1:200 |

| | | | | | |
|---|---|---|---|---|---|
| Anti-CCNO antibody | Rabbit | WB | Atlas | HPA050090 | 1:500 |
| Anti-Nanog antibody | Goat | IF | R&D systems | AF1997 | 1:100 |
| Anti-AFP antibody | Mouse | IF | Sigma-Aldrich | A8452 | 1:1000 |
| Anti-SMA antibody | Mouse | IF | Pierce | MA1-12772 | 1:1000 |
| Anti-TUJ1 antibody | Mouse | IF | Covance | MMS-435P | 1:1000 |
| Anti-HA antibody | Rabbit | WB, IF | Cell Signaling | CST 3724 | 1:1000 |
| Anti-β-tubulin antibody | Mouse | WB | Proteintech | 66240-1-Ig | 1:1000 |
| Anti-RFX2 antibody | Rabbit | IF | Sigma-Aldrich | HPA048969 | 1:250 |
| Anti-RFX3 antibody | Rabbit | IF | Sigma-Aldrich | HPA035689 | 1:250 |

**Table S3.** Genotypes and phenotypes of patients with *CCNO* pathogenic variations

| Individual (Ancestry) | *CCNO* Genotype | Known Consanguinity | Gender | Airway Disease | nNO |
|---|---|---|---|---|---|
| **F**, 19GM00243 (North Africa) | c.248_252dup p.(Gly85Cysfs*11) homozygous | Yes | Male | NNRD<br>Bronchitis<br>Bronchiectasis<br>Rhinosinusitis<br>Otitis media<br>Mild developmental delay | Low§ |
| **G**, 19GM00351 (France) | c.337G>A p.(Glu113Lys) c.592A>T p.(Lys198*) compound heterozygous | No | Male | NNRD<br>Bronchitis<br>Bronchiectasis<br>Rhinosinusitis<br>Otitis media<br>Inborn deafness | Low§ |
| **H**, 18GM01179 (France) | c.248_252dup p.(Gly85Cysfs*11) c.793dup p.(Val265Glyfs*106) compound heterozygous | No | Male | NNRD<br>Bronchitis<br>Bronchiectasis<br>Rhinosinusitis<br>Otitis media | Low§ |
| **I**, 19GM01068 (Turkey) | c.258_262dup p.(Gln88Argfs*8) homozygous | Yes | Male | NNRD<br>Bronchitis<br>Rhinosinusitis<br>Otitis media | NA |
| **J**, OP-1246 II1 (Germany) | c.262_263insGGCCC p.(Gln88Argfs*7) homozygous | Yes | Male | NNRD<br>recurrent pneumonia<br>Bronchitis<br>Rhinitis<br>Cough | NA |
| **K**, OP-642 II1 (Germany) | c.926delC p.(Pro309Argfs*17) homozygous | Yes | Female | NNRD<br>Otitis media<br>Sinusitis<br>recurrent pneumonia<br>bronchitis | Low§ |
| **L**, OI-66 II1 (Israel) | c.638T>C; p.(Leu213Pro) homozygous | Yes | Male | Bronchiectasis<br>recurrent pneumonia<br>Otitis<br>Sinusitis<br>Cough | NA |
| **M**, OI-104 II1 (Israel) | c.262_263insGGCCC p.(Gln88Argfs*7) homozygous | NA | Male | Bronchiectasis<br>recurrent pneumonia<br>Cough | NA |
| **N**, OP-151 II1 (Austria) | c.262_263insGGCCC p.(Gln88Argfs*7) homozygous | No | Male | NNRD<br>recurrent pneumonia<br>Rhinitis<br>Cough | Low§ |
| **O**, OP-971 II1 (Germany) | c.716A>G p.(His239Arg) homozygous | Yes | Female | NNRD<br>recurrent pneumonia<br>Bronchiectasis<br>Otitis media | NA |

| | | | | | |
|---|---|---|---|---|---|
| **P**, OP-1367 II1 (Germany) | c.262_263insGGCCC p.(Gln88Argfs*7) homozygous | Yes | Male | NNRD<br>recurrent pneumonia<br>Rhinosinusitis<br>Bronchiectasis<br>Otits media<br>Cough | NA |
| **Q**, OP-1777 I5 (Kuwait) | c.252_253insTGCCC p.(Gly85Cysfs*10) homozygous | NA | Female | NA | NA |
| **R**, OP-1777 II1 (Kuwait) | c.252_253insTGCCC p.(Gly85Cysfs*10) homozygous | NA | Male | NA | NA |
| **S**, OP-1777 II2 (Kuwait) | c.252_253insTGCCC p.(Gly85Cysfs*10) | NA | Female | NA | NA |
| **T**, OP-1977 II1 (Germany) | c.262_263dupGGCCC p.(Gln88Argfs*7) homozygous | NA | M | Middle lobe atelectasis<br>Bronchiectasis<br>Rhinitis<br>Cough | Low§ |

§: <77nL/min; nNO: nasal nitric oxide; TEM: transmission electron microscopy; NNRD: neonatal respiratory distress; NA: not assessed

**Table S4.** Genotypes and phenotypes of control PCD patients

| **Individuals (Ancestry)** | **Gene Genotype** | **Known Consanguinity** | **Gender** | **Laterality defect** | **Airway Disease** | **nNO** | **HSVA** (CBF) | **TEM defects** |
|---|---|---|---|---|---|---|---|---|
| **A**, 18GM01423 (South Asia) | *DNAH5:* c.841_842insCTTCCGC p.(Val281Alafs*20) homozygous | Yes | Female | No | NNRD Bronchitis Bronchiectasis Rhinosinusitis Nasal polyposis Otitis media | Low§ | Immotile cilia | ODA |
| **B**, 20GM000711 (France) | *DNAH5:* c.2710G>T p.(Glu904*) c.9897+1del p.? compound heterozygous | No | Female | Yes | NNRD Bronchitis Bronchiectasis Rhinosinusitis Otitis media | Low§ | NA | ODA & IDA |
| **C**, 22GM000288 (France) | *CCDC39:* c.2347_2351delTTTCA p.(Phe783Thrfs*3) homozygous | No | Male | No | NNRD Bronchitis Bronchiectasis Rhinosinusitis Otitis media | Low§ | Dyskinetic and immotile cilia | IDA with MTD |
| **D**, 20GM000324 (Sub-Saharan Africa) | *CCDC39:* c.876del p.(Lys292Asnfs*16) c.2158+5del p.? compound heterozygous | No | Male | No | Bronchitis Bronchiectasis Rhinosinusitis Serous otitis | Low§ | Dyskinetic cilia | IDA with MTD |
| **E**, 18GM00086 (Caribbean) | *DNAH11:* c.11182dup p.(Leu3728Profs*10) homozygous | No | Female | Yes | Bronchitis Bronchiectasis Rhinosinusitis Serous otitis | Low§ | Immotile cilia | Normal ultrastructure |

§: <77nL/min; nNO: nasal nitric oxide; TEM: transmission electron microscopy; NNRD: neonatal respiratory distress; NR: non relevant (child); NA: not assessed; ODA: outer dynein arm defect; IDA: inner dynein arm defect; MTD: microtubular disorganization

### Materials and methods

#### Transgenic mice

All animal studies were performed in accordance with the guidelines of the European Community and French Ministry of Agriculture and were approved by the "Direction départementale de la protection des populations de Paris" (Approval number Ce5/2012/107; APAFiS #9343). Most of the mouse strains have already been described: *Ccno*$^{RA}$ are from Sebastian J. Arnold's laboratory (Funk et al., 2015) and *Ccno*$^{KO}$ are from Gabriel Gil-Gómez's laboratory (Núnez-Ollé et al., 2017). Wild-type are either RjOrl:SWISS (Janvier labs), or the WT and/or HT pups from the same litters as the different *Ccno* mutants.

#### Primary brain ependymal cell cultures and transfections

The ependymal culture has been previously described (Delgehyr et al., 2015). Briefly, newborn mice (P0-P2) were sacrificed by decapitation. The brains were dissected in Hank's solution (10% HBSS, 5% Hepes, 5% sodium bicarbonate, 1% Penicillin/Streptomycin (P/S) in pure water) and the telencephalon were cut manually into pieces, followed by enzymatic digestion (DMEM glutamax, 33% papain (Worthington 3126), 17% DNAse at 10 mg/ml, 42% cystein at 12 mg/ml) for 45 minutes at 37°C in a humidified 5% CO2 incubator. Digestion was stopped by addition of a solution of trypsin inhibitors (Leibovitz Medium L15, 10% ovomucoid at 1 mg/ml, 2% DNAse at 10 mg/ml). The cells were then washed in L15 and resuspended in DMEM glutamax supplemented with 10% fetal bovine serum (FBS) and 1% P/S in a Poly-L-lysine (PLL)-coated flask. Ependymal progenitors proliferated for 4-5 days until confluence before shaking (250rpm) overnight. Pure confluent astroglial monolayers were replated at a density of $7{,}5\times10^3$ cells/µL (corresponding to days in vitro "DIV" -1) in DMEM glutamax, 10% FBS, 1% P/S on PLL coated coverslides for immunofluorescence experiments and maintained overnight. The medium was then replaced by serum-free DMEM glutamax 1% P/S, to trigger ependymal differentiation in vitro (DIV 0). Transfections were performed in suspension at DIV -1 using the Jetprime (Polyplus) system. Plasmids used are pCAG-H2B-GFP from Xavier Morin's laboratory (Hadjantonakis and Papaioannou, 2004), pCAGGS-Mcidas from Stavros Taraviras' laboratory (Kyrousi et al., 2015) and pcDNA3myc-hCyclinO from Gabriel Gil-Gómez's laboratory.

#### Immunostainings

Brain lateral ventricles and tracheae samples were permeabilized in PBS 1X with 0,1% and 0,5% Triton X-100 solutions, respectively, for 1 minute at RT, then fixed in methanol at -20°C for 7 minutes. Cell cultures were fixed in methanol at -20°C for 10 minutes or PFA 4% at RT for 10 minutes. Tissues and cells were preblocked in PBS 1X with 0.2% Triton X-100 and 10% FBS (blocking solution) before incubation with primary then secondary antibodies. All these were incubated overnight at 4°C or for 1h at RT in the primary antibodies diluted in blocking solution. Nuclei were counterstained with a 1:1500 Hoechst solution (33342, from a 20 mg/ml stock, Sigma-Aldrich), containing the secondary antibodies for 1h at RT. Primary antibodies used are listed in **Supplementary Table S2** and secondary antibodies are species-specific Alexa Fluor® secondary antibodies (1:400; Invitrogen). Finally, the wholemounts were redissected to keep only the thin lateral walls of the LV (Delgehyr et al., 2015) which were mounted with Fluoromount-G mounting medium (Southern Biotech,

0100-01). Cell cultures on coverslips were mounted on glass slides with Fluoromount-G. Fluoromount-mounted slides were stored at 4°C.

### Microscopy

#### *Epifluorescence microscopy*

Fixed cells were examined with an upright epifluorescence microscope (Zeiss Axio Observer.Z1) equipped with an Apochromat ×63 (NA 1.4) oil-immersion objective and a Zeiss Apotome with an H/D grid. Images were acquired using Zen with 500-nm z-steps. When better resolution was needed, confocal image stacks were collected with a 63 x/1.4 Oil objective on an inverted LSM 880 Airyscan Zeiss microscope with 440, 515, 560 and 633 laser lines with 180 or 250-nm z-steps and Zen2 software with Airyscan mode Z-stack projections are shown for all epifluorescence microscopy images.

#### *Transmission electron microscopy*

Cultured cells were fixed in 2.5% glutaraldehyde and 4% PFA, treated with 1% OsO4, washed and progressively dehydrated. The samples were then incubated in 1% uranyl acetate in 70% methanol, before final dehydration, pre-impregnation with ethanol/epon (2/1, 1/1, 1/2) and impregnation with epon resin. After mounting in epon blocks for 48 h at 60°C to ensure polymerization, ultra-thin sections (70 nm) were cut on an ultramicrotome (Ultracut E, Leica) and analysed using a Philips Technai 12 transmission electron microscope.

### Single-cell RNA-seq of in vitro differentiating multiciliated cells

Ependymal cells were cultured in flask. After 2 days of in vitro differentiation, flasks were rinsed with PBS 1X twice and treated by enzymatic cell dissociation (Trypsin, 1mL) for 10 min and trituration to obtain a single cell suspension. Digestion was stopped by addition of 1 ml of fetal bovine serum (FBS). The cells were then washed in serum-free DMEM glutamax 1% P/S and resuspended in HBSS-0.1% BSA for single-cell RNA-seq library preparation. Cell cultures from three different animals with the same genetic background and treatment were pooled together; to multiplex them, samples were labelled using a cell surface protein labelling strategy following manufacturer's instructions (https://asets.ctfa-sets.net/an68im79xiti/5KA1NbZdTOam8A0yq6KyC2/f3e0479ff7b1c1633e6ddf8959a88a3bCG000149_DemonstratedProtocol_CellSurfaceProteinLabeling_Rev_A.pdf), using TotalSeq® hashtag antibodies (BioLegend). However, the efficiency of the cell hashing did not allow to unambiguously separate cells coming from different animals and was therefore not directly used in downstream analysis. Cell suspensions were passed through a 40 µm Flowmi cell strainer (BelArt) and cell concentrations were carefully evaluated with a Countess FL automated cell counter (Thermofisher). For single cell RNA-seq, cells were partitioned with a Chromium equipment (10X Genomics) and libraries were prepared using the standard Single Cell 3' v3.1 protocol (10X Genomics). Sequencing was performed on an Illumina NextSeq 500 sequencing machine following manufacturer's instructions, on a high Flowcell, using the following sequencing cycles: 28b for read 1, 55b for read 2, 8b for index.

### Computational analysis of single-cell RNA-seq data

#### *Pre-processing of scRNAseq of in vitro differentiating multiciliated Ccno$^{KO}$ cells*

Fastq file demultiplexing, barcode processing, gene counting, and aggregation were made using the Cell Ranger software following 10X Genomics guidelines. 10X Genomics mm10 genome reference and gene annotations compiled in 2020 were used (https://support.10xgenomics.com/single-cell-gene-expression/software/release-notes/build#mm10_2020A). Empty cells (detected by emtpyDrops from DropletUtils) were filtered out and only protein-coding genes were retained.

#### *Annotation transfer and UMAP projection between WT and Ccno$^{KO}$ scRNAseq datasets*

scmap package (Kiselev et al., 2018) was used to transfer cluster annotations from WT to Ccno$^{KO}$ cells. For visualization, projection of the *Ccno$^{KO}$* cells onto the WT UMAP embedding (Serizay et al.) was performed using Seurat (**Fig. S4A**) (Hao et al., 2021).

#### *Merging WT or Ccno$^{KO}$ scRNAseq datasets*

Genotype correction and merging of WT (Serizay et al.) and Ccno$^{KO}$ single-cell RNA-seq datasets was performed using fast MNN correction from batchelor package (Haghverdi et al., 2018), using genes found enriched (fold-change > 2, adjusted p-value <= 0.05) in any of the annotated/transferred clusters with scran package (Lun et al., 2016).

#### *Cell cycle phase annotation of merged WT or Ccno$^{KO}$ cells*

Putative cell cycle phases were transferred from a recent scRNAseq dataset of proliferating neural stem cells with annotated cell cycle phases (O'Connor et al., 2021) using SingleR (Aran et al., 2019). G0, G1 and Late G1 labels were collapsed to a single "G0/G1" label.

#### *Trajectory analysis of merged WT or Ccno$^{KO}$ cells*

Trajectory and pseudotime inferences were performed using the shared MNN-corrected PCA embedding of merged WT and Ccno$^{KO}$ cells (**Fig. S4B**) with slingshot (Street et al., 2018), specifying start (proliferating progenitors) and end clusters (terminally differentiated MCC) to orientate the trajectory. Computed pseudotime was scaled between 0 and 1 for clarity. Continuous gene expression along the slingshot-inferred trajectory was modelled using a generalized additive model.

#### *Characterization of primordial deuterosomal cells in WT or Ccno$^{KO}$ scRNAseq datasets*

The pseudotime value containing 95% of all Ccno$^{KO}$ cells was determined, and subsequently used to split the original "Early Deuterosomal Cells" cluster (**Fig. S4A**) into "Primordial deuterosomal" and "Early deuterosomal" cells (**Fig. 3A, Fig. S4C**).

Ccno$^{KO}$ vs. WT differential gene expression analysis was performed specifically on cells from the "Primordial deuterosomal" cluster. To control for technical variations between individuals and/or replicates, we first phased each cell in each dataset (3 individuals pooled for each of the 2 WT replicates, and 3 individuals pooled in the

Ccno$^{KO}$ dataset) using scSplit (Xu et al., 2019). I then used the phasing and the replicate annotations to group WT (Ccno$^{KO}$) primordial deuterosomal into 6 (3) meta-cells using the ` aggregateAcrossCells ` function from the scuttle package. Each meta-cell thus contains cells from a separate individual, with a specific genetic background. I then performed a differential expression analysis on these pseudo-bulk meta-cells using DESeq2 (Love et al., 2014), treating the replicate and genetic background as confounding variables. Genes differentially expressed between Ccno$^{KO}$ and WT primordial deuterosomal cells (absolute fold-change > 2, adjusted p-value <= 0.1) were finally recovered. GSEA was performed with clusterProfiler (Wu et al., 2021).

### Generation of human epithelial airway cells from human embryonic stem cell (hESC)

#### *Human embryonic stem cell (hESC) culture*

H9 hESCs were routinely cultured on vitronectin-coated tissue culture dishes in Essential 8 (E8) medium (Life Technologies) in an incubator with 5% CO2, at 37°C.

#### *Construction of CRISPR sgRNA vectors and donor template*

Three CRISPR sgRNAs, targeting exons 1 and 2, of the human *CCNO* gene were designed using CHOPCHOP (https://chopchop.cbu.uib.no/) and cloned into the pHF1-Cas9 plasmid. The sgRNA sequences are provided in Table S1.

#### *Nucleofection*

H9 cells were dissociated into single cells with Accutase (Stem Cell Technologies). Two million cells were mixed with 10 µg pHF1-Cas9-sgRNA and nucleofected using the Lonza Amaxa 4D Nucleofector (Lonza). E8 medium supplemented with 5.25 µg/ml Blasticidin was applied to the cells 24 hours post-nucleofection. Cells were fed E8 media every day until colonies were large enough for picking and genotyping.

#### *Genotyping*

To extract genomic DNA, picked colonies were incubated for 55ºC for 1 hour followed by a 5 min incubation at 95ºC in a 20 µl reaction containing 1X detergent (0.05% IGEPAL CA630, 0.05% Tween-20), proteinase K and 1X TE buffer. 1 µl of genomic DNA was used in a PCR reaction containing 1X Primestar Max Mastermix (Takara) and genotyping primers. The thermal cycling conditions used were as follows: 10 s at 98 °C, followed by 35 cycles of 10 s at 98 °C, 5 s at 55 °C and 1 min at 72 °C. The PCR products were then sent to Bio Basic Asia Pacific Pte Ltd for sequencing. Genotyping primer sequences are listed in Table S1.

#### *Clonal expansion*

H9 clones with frame-shift mutations were dissociated into single cells using Accutase and seeded onto vitronectin-coated plates in E8 medium supplemented with CloneR (Stem Cell Technologies). Cells were fed with fresh E8 medium daily until colonies grew large enough for picking and genotyping. Clones with homozygous frame-shift mutations were expanded and frozen stocks were made using Cryostor (Stem Cell Technologies).

***Construction of wild-type (WT) and mutant CCNO expression vectors***

WT and mutant CCNO cDNA (c.431_444del) were cloned into the pCS2+ expression vector. FLAG tags were added to the N termini and HA tags were added to the C termini of both WT and mutant sequences. Primer sequences are provided in Table S1.

***Transfection of HEK293T cells***

500,000 HEK293T cells were seeded onto wells of a 6-well plate in MEF media and allowed to attach overnight in an incubator at 37ºC. The following day, cells were transfected with 1 μg pCS2+CCNO WT, pCS2+CCNO c.431_444del or pCS2+eGFP using Lipofectamine 2000 (Life technologies). Cells were harvested for Western blotting or immunofluorescence 48 hours-post transfection.

***Trilineage differentiation***

H9 cells were dissociated into small clumps using Accutase (Stem Cell Technologies) and collected in a falcon tube. Cells were collected into a pellet by centrifugation at 1200 rpm for 5 minutes. The cell pellet was resuspended in MEF medium (15% FBS, 1% Glutamax in DMEM F12 Advanced), and the resulting cell suspension transferred into low adhesion tissue culture dishes to form embryoid bodies. The medium was changed every other day for 7 days. The embryoid bodies formed were collected and plated onto 0.1% gelatin-coated tissue culture dishes. Cellular outgrowths were fed every other day with fresh MEF medium for a further 7 days before they were fixed with 4% paraformaldehyde for analysis by immunofluorescence.

***Karyotype analysis***

H9 CCNO subclones, 52-3 and 20-1 cultured to 70-80% confluency, were sent to the KK Women's and Children's Hospital (Singapore) Cytogenetics department for karyotype analyses.

***Immunofluorescence***

Cells were washed with DPBS and then fixed in 4% PFA for 20 min at RT. PFA was removed and cells were washed with DPBS before they were blocked and permeabilized in blocking buffer (10% donkey serum, 0.1% Triton X-100 in DPBS) for 1h at RT. Cells were then incubated with primary antibodies diluted in staining buffer (1% donkey serum, 0.1% Triton X-100 in DPBS) overnight at 4°C. Excess primary antibodies were removed and the cells were washed 3 times with DPBS before incubation with fluorescence-conjugated secondary antibodies diluted in staining buffer for 1 h in the dark at RT. Cells were washed 3 times with DPBS to remove excess and unbound antibodies. NucBlue Fixed Cell ReadyProbes Reagent (DAPI, ThermoFisher Scientific) diluted in DPBS at 1 drop/ml was then added to each well. Cells were imaged using the Leica FV-3000. ImageJ was used for image processing and analysis. Primary antibodies used can be found in **Supplementary Table 2**. Secondary antibodies are species-specific Alexa Fluor® secondary antibodies (1:1000; Invitrogen).

***RNA extraction, cDNA synthesis and quantitative real-time PCR (qPCR)***

Qiagen RNeasy kit was used to extract total RNA from cells. 500 ng of purified RNA was converted into cDNA using the High-Capacity cDNA Reverse Transcription Kit (Applied Biosystems). QPCR was performed using the QuantStudio 7 Flex Real-Time PCR System with samples run in duplicates and normalized to housekeeping gene *ACTB*. Primer sequences are listed in **Table S1**. Adult lung mRNA used as a positive control for QPCR experiments was obtained from Biochain (Total RNA – Human Adult Normal Tissue: Lung lower left lobe; Cat #R1234152-50; Lot #B6050780).

***Protein extraction and western blot***

Whole cell lysates were prepared by lysing cells in Pierce RIPA buffer (ThermoFisher Scientific) supplemented with complete protease inhibitor cocktail (Calbiochem). 10 µg protein was resolved on a 7.5% SDS PAGE gel and then transferred onto PVDF membranes via semi-dry transfer method (Bio-rad). Membranes were blocked in 5% skim milk in Tris-buffered saline (TBST: 0.05 M Tris, 0.138 M NaCl, 0.0027 M KCl, pH 8.0) with 0.1% Tween-20 (Sigma-Aldrich) for 1h at RT. Membranes were then incubated with primary antibodies at 4 °C overnight. Membranes were then washed 3 times in TBST and then incubated with HRP-linked secondary antibodies for 1h at RT. Membranes were washed 3 times with TBST before proteins were visualized using the SuperSignal West Femto Maximum Sensitivity Substrate (ThermoFisher Scientific) on the Chemidoc (Bio-rad). Primary and secondary antibodies are listed in Table S2.

***Differentiation of hESCs into lung progenitors***

H9 cells were dissociated into single cells with Accutase for 5 min at 37°C. 400,000 cells were seeded into each well of a 12-well plate and left in the incubator to attach overnight. Cells were fed fresh E8 media the next day (Day -1, D-1). At D0, H9 cells were differentiated into definitive endoderm (DE) cells using a protocol previously published by Vallier *et al.,* 2009. Briefly, on D0, cells were treated 100 ng/ml Activin A, 80 ng/ml FGF2, 10 µM LY294002, 3 µM CHIR99021, 10 ng/ml BMP4 in CDM-PVA medium. On D1, cells were treated with 100 ng/ml Activin A, 80 ng/ml FGF2, 10 µM LY294002 and 10 ng/ml BMP4 in CDM-PVA medium. On D2, cells were treated with 100 ng/ml Activin A, 80 ng/ml FGF2, 1X B27, 1X NEAA in RPMI medium. To differentiate the DE cells into anterior foregut and subsequently lung progenitors, the protocol published by McCauley *et al.,* 2017 was used. To drive DE cells towards anterior foregut formation, cells were treated with 10 µM SB431542 and 2 µM Dorsomorphin (DSM) in basal medium (1X B27, 1X N2, 1X Glutamax, 1 mM HEPES in DMEM F12 Advanced) for 3 days. To differentiate AFE cells into lung progenitors, cells were treated with 3 µM CHIR, 10 ng/ml BMP4, 10 ng/ml FGF7, 10 ng/ml FGF10 and 50 nM RA in basal medium for 9 days.

***Fluorescence-activated cell sorting (FACS)***

For sorting at Day 15, differentiated H9 cells were washed once with DPBS and then dissociated into single cells with TryPLE express. TrypLE express was diluted out with DMEM Advanced F12 and removed by centrifugation at 1200 rpm for 5 min. The cell pellet was resuspended in FACS buffer supplemented with 2% penicillin/streptomycin and 10 µM Y-27632. The cell suspension was passed through a 40 µm cell strainer to remove cell clumps before cells were counted using the Scepter Cell Counter (Merck Millipore). Cell concentration was adjusted to 1 million cells/100 µl FACS buffer. 0.5 µl isotype control or 0.5 µl Human AL-CAM/CD166 Phycoerythrin (PE)-conjugated antibody (Clone 105902) (R&D Systems) was added per 100 µl. Cells were stained for 30 minutes in the dark at 4°C. Excess antibodies were diluted out and removed by adding FACS buffer followed by centrifugation at 1200 rpm for 5 minutes. Cell pellets were resuspended in 500 µl FACS buffer. Samples were FACS sorted with the unstained and isotype-stained H9-derived lung progenitors as a negative control for the gating parameters.

For sorting at Day 25-30 to collect airway basal cells, lung organoids embedded in Matrigel were washed with DPBS once. Organoids were then incubated with TrypLE express for 30 min at 37°C. DMEM F12 advanced was used to dilute out TrypLE express and then cells were collected into a pellet by centrifugation at 1200 rpm for 5 min. The cells were resuspended in FACS buffer supplemented with 2% penicillin/streptomycin and 10 µM Y-27632, passed through a 40 µm cell strainer. Cells were counted and cell concentration was adjusted to 1 million cells/ 100 µl FACS buffer. 0.5 µl of each isotype control or 0.5 µl Human NGFR (APC)-conjugated antibody and 0.5 µl EpCAM (PE)-conjugated antibody were added per 100 µl. Cells were stained for 30 minutes in the dark at 4°C. Excess antibodies were diluted out and removed by adding FACS buffer followed by centrifugation at 1200 rpm for 5 minutes.

Cell pellets were resuspended in 500 µl FACS buffer prior to FACS sorting. Unstained and isotype-stained H9-derived lung cells were used as negative controls for the gating parameters.

***Proximal lung organoid culture***

To obtain proximal lung organoids, a protocol published by Hawkins et al., 2021 was used. Briefly, Day 15-sorted CD166+ cells were resuspended in basal medium containing 250 ng/ml FGF2, 100 ng/ml FGF10, 50 nM Dexamethasone (Sigma-Aldrich), 0.1 mM 8-Bromoadenosine 30,50-cyclic monophosphate sodium salt (cAMP, Sigma-Aldrich) and 0.1 mM 3-Isobutyl-1-methylxanthine (IBMX) (Sigma-Aldrich). Undiluted growth factor-reduced Matrigel (Corning) was added to the cell suspension at a 1:1 ratio after which 40 µl of this cell suspension was added to the middle of each 24-well plate to create a Matrigel drop. The plates were then returned to the incubator to allow the Matrigel drops to solidify for at least 30 min before medium is overlaid over the drops. Y-27632 was included in the medium for the first 24 h to improve cell survival. Medium was refreshed every other day for ~2 weeks before proximal lung organoids were harvested for cell sorting.

***Terminal differentiation of ESC-derived basal cells cells***

150,000 cells were seeded onto 6.5 mm Matrigel-coated Transwell inserts. When cells reached 100% confluence, media was removed from both chambers and PneumaCULT ALI medium (Stem Cell Technologies) was added only to the bottom chamber. Medium was refreshed every other day for 2 weeks before cells were harvested for mRNA or fixed in 4% PFA or methanol for immunostaining.

***Western blotting to detect endogenous CCNO from ALI cultured hESCs***

6 transwells of wild type or CCNO mutant cells, respectively, were lysed with 300 µl RIPA buffer (Thermo Fisher Scientific). The cell lysates were sonicated for 10 seconds and spun down for 15 minutes at 12000 *g*. 200 µl cell lysate was boiled with 200 µl SDS loading buffer. Subsequently, the cell lysates were separated on SDS-PAGE gels, transferred to PVDF membrane and incubated with blocking buffer (3% BSA, 0.1% tween in PBS). Previously validated rabbit anti-*CCNO* antibody (Atlas HPA050090, validated in Wallmeier et al. (2014) Nature Genetics 46:646-651) was used for detection of endogenous CCNO protein.

***Immunofluorescence staining of ALI cultured hESCs and microscopy***

Cells grown on transwells were fixed with ice cold methanol at -20ºC for 15 min. Cells were then blocked with blocking buffer (3% bovine serum albumin in PBS) for 1 hour, followed by 1 hour incubation with relevant primary antibodies (listed in **Supplementary Table S2**) at room temperature. After three washes with wash buffer (0.1% triton in PBS), cells were incubated with secondary antibodies and DAPI for 1 hour. After three washes with wash buffer, the stained cells were mounted on glass slides with fluorescence-mounting medium and imaged with an Olympus FluoView upright laser confocal microscope.

**Transmission electron microscopy on human respiratory cells**

***Patients A to I***

Samples of airway epithelial cells were obtained from nasal or bronchial biopsies of patients. They were fixed by 2.5% glutaraldehyde in 0.045 M cacodylate buffer, pH 7.4, for a period of two hours at a temperature of 4°C. Subsequently, the samples were postfixed with osmium tetroxide ($OsO_4$) and processed for transmission electron microscopy in accordance with standard procedures. The ultrathin sections were examined at different magnification of 6,000 to 100,000).

***Patients J to T***

Respiratory epithelial cells were obtained by nasal brush biopsies from the middle turbinate with nasal biopsy brushes (Engelbrecht Medicine and Laboratory technology). For transmission electron microscopy analyses, airway cells were directly suspended in 2.5% glutaraldehyde and were fixed overnight at 4°C. Afterwards, cells were stained with 1% osmium tetroxide. For complete permeabilization, the stained material was then incubated overnight at 4°C with a 1,2-epoxypropane-epone mixture in a 1:1 ratio. The samples was then embedded in epon, before they were transferred onto cupper grids. Contrasted with uranyl acetate and Reynold's lead citrate sample material was finally visualized using the Philips CM10 microscope.

### Quantification and statistical analysis

Images used for intensity quantification were all taken with the same microscopy parameters. Nuclear and cytoplasmic intensity was measured used an in-house built ImageJ macro. Hoechst staining was segmented using morphological filtering and the segmented nucleus area was used to measure the mean fluorescence intensity of the channel of interest per nucleus. Cytoplasmic fluorescence intensity quantifications were performed by manual outlining of the cellular fluorescence signal. Nuclear and cytoplasmic measurements were then classified based on differentiation stages and divided by the mean value per coverslip of the centrosome stage signal and normalized as that y=1 corresponds to the mean centrosome stage intensity.

All graphs and statistical analyses were obtained using GraphPad Prism software. Data were obtained from at least three independent experiments unless differently stated.
